# Specificity-driven protein binder design with Odin-Multi

**DOI:** 10.64898/2026.09.08.749745

**Authors:** Valentas Brasas, Charlotte R. Christensen, Kasper H. Björnsson, Victor Emil Møller, Beatrice Scapolo, Melisa Benard-Valle, Mads Mørup Nygaard, Jens Christian Nielsen, Sine Reker Hadrup, Kristoffer H. Johansen, Timothy P. Jenkins

**Author notes:** These authors contributed equally (Valentas Brasas, Charlotte R. Christensen, Kasper H. Björnsson, Victor Emil Møller).

## Abstract

A useful protein binder is defined as much by what it does not bind as by what it does. Some applications call for one binder to cover a family of related targets; others require it to distinguish a single member from near-identical relatives. Yet, widely used deep-learning-based *de novo* design methods typically optimise one interaction at a time, leaving cross-reactivity and specificity to emerge during downstream screening. Here we present Odin-Multi, a binder design framework that optimises a shared binder sequence against several complexes simultaneously, applying attractive objectives to on-targets and repulsive objectives to off-targets.

We benchmarked Odin-Multi *in silico* across three systems representing distinct cross-reactivity and specificity challenges: class B1 G protein-coupled receptors (GPCRs), testing cross-reactivity across multiple therapeutically relevant receptors; short-chain three-finger toxins, testing cross-reactivity across homologous toxin family members; and peptide-MHC (pMHC) complexes, testing specificity between near-identical target and off-target surfaces. For pairs of related class B1 GPCRs, 83.5 to 96.8% of jointly optimised designs exceeded an interaction-confidence threshold for both targets, compared with 6.8 to 36.3% of designs from single-target campaigns. For two short-chain three-finger neurotoxins, 9.2% of jointly optimised designs exceeded the corresponding threshold for both targets, compared with 0.8% of designs optimised against one toxin alone. Finally, in a pMHC specificity benchmark where target and off-target differed only in a single peptide residue, counter-selection increased the fraction of designs satisfying both the target-confidence criterion and a target-to-off-target interaction-confidence ratio of 2.5 from 6.0% to 14.2%.

Experimental screening produced leads consistent with both design regimes in the two systems tested *in vitro*. We identified a cross-reactive toxin minibinder showing apparent nanomolar binding to the neurotoxin Erabutoxin A and to a candidate NK-shNTx-containing fraction from *Naja kaouthia* venom (higher-affinity fitted components of 11.95 and 34.43 nM, respectively), and a pMHC minibinder with greater target-to-off-target discrimination than a previously reported design. By treating cross-reactivity and specificity as explicit design objectives rather than screening outcomes, Odin-Multi widens the range of binding behaviours accessible to computational design.

## 1 Introduction

In therapeutic, diagnostic, and research applications, protein binders must often discriminate between closely related molecular surfaces, engaging on-targets while sparing homologous, structurally similar, and unrelated off-targets; or in some settings, engaging an entire family of related proteins while sparing the rest of the proteome [1, 2, 3]. Indeed, off-target receptor engagement can drive severe toxicity in T-cell receptor (TCR) therapies [4], while narrow species coverage limits the utility of monoclonal antivenoms relative to polyclonal sera [5, 6, 7].

While recent advances in deep learning have transformed *de novo* binder design [8, 9, 10, 11, 12, 13, 14, 15] and hallucination as well as diffusion methods can now generate binders against a wide range of targets with increasingly high experimental success rates [9, 15], they currently typically optimise a single binding interaction. As such, specificity and breadth of targets bound are only addressed through binding-site selection or downstream through structural filtering [13, 16, 17], *in silico* cross-panning [18] or lab-based screening [19]. Such selection can identify candidates with the desired interaction profile when these already occur among the designs produced, but does not influence which designs arise. This limitation is particularly problematic when on-targets and off-targets share structural features, or when broad recognition of a conserved feature is required [1, 20]. Explicit multi-target optimisation therefore provides a principled means of incorporating specificity or cross-reactivity objectives during candidate generation [21, 22, 23, 24].

Here we introduce Odin-Multi, a binder design framework that incorporates multi-target and multi-off-target interaction profiles directly into the optimisation objective. Within each design trajectory, a single binder sequence is evaluated against a user-defined set of on-targets and, optionally, off-targets through parallel structure prediction, producing a separate objective for each complex. On-target objectives promote binding, whereas off-target objectives discourage it, allowing all specified interaction contexts to shape the same sequence during optimisation. Odin-Multi thereby extends binder hallucination beyond optimising whether a binder engages a single target, towards shaping how the same sequence interacts across multiple on-targets and competing off-targets [15]. Depending on the specified contexts, this formulation can encode specific, crossreactive, or selectively cross-reactive interaction profiles.

We benchmarked Odin-Multi computationally across three biological systems spanning two design modes: cross-reactivity, where a single binder is designed to engage several related on-targets, and specificity, where a binder is designed to engage an on-target while avoiding one or more defined off-targets. We then validated one system from each mode experimentally. As a first computational cross-reactivity benchmark, we targeted the closely related and therapeutically relevant class B1 G protein-coupled receptors (GPCRs) GLP-1R, GCGR, and GIPR [25, 26], testing whether a single peptide sequence could achieve balanced predicted interactions across multiple related receptors. As a second cross-reactivity case, we computationally benchmarked and experimentally validated targeted cross-reactivity against short-chain three-finger toxins from elapid snake venoms, where broad neutralisation across the toxin family is key to therapeutic efficacy [5, 20, 27, 28]. Finally, we benchmarked and experimentally validated exclusion-based specificity against human leukocyte antigen (HLA) presenting the cancer-testis antigen NY-ESO-1, SLLMWITQC, a setting in which the on- and off-targets share an identical scaffold and differ only in the presented peptide, representing one of the most stringent discrimination problems in current protein design [18, 16, 29]. Across these systems, Odin-Multi enriched predicted cross-reactivity in the GPCR and three-finger toxin *in silico* benchmarks and promoted predicted specificity in the peptide–MHC (pMHC) *in silico* benchmark. Experimental testing identified a cross-reactive three-finger toxin minibinder and a *de novo* pMHC minibinder showing greater on-target/off-target discrimination than the previously reported prototype [18].

By incorporating specificity and cross-reactivity directly into the design objective, Odin-Multi extends current *de novo* binder design beyond single-interaction optimisation and provides a general framework for generating candidates with interaction profiles tailored to therapeutic, diagnostic, and biotechnological applications.

## 2 Results

### 2.1 Odin-Multi enables multi-context binder optimisation

Odin-Multi extends AlphaFold2-based binder hallucination from a single-complex objective to a unified multi-complex optimisation framework (Fig. **1**). Within each design trajectory, the same binder sequence is evaluated independently against all specified on-target and off-target structures, producing separate structure predictions and objectives for each complex. These context-specific objectives jointly update the shared sequence, allowing multiple desired and undesired interactions to influence optimisation within the same trajectory. Sequence optimisation used a three-stage protocol in which the sequence was parameterised in logit space, while the representation supplied to AlphaFold2 progressed from continuous logits, through temperature-annealed softmax probabilities, to straight-through one-hot sequences. Campaign-specific iteration counts and subsequent discrete refinement steps are described in Methods. Campaign-specific LigandMPNN [30] or ProteinMPNN [10] losses were optionally used to encourage sequence–structure compatibility. We then evaluated Odin-Multi computationally in three benchmark settings: two cross-reactivity cases, in which a single sequence was optimised to bind multiple on-targets, and one specificity case, in which binding to an on-target was optimised while binding to defined off-targets was penalised.

**Figure 1:**
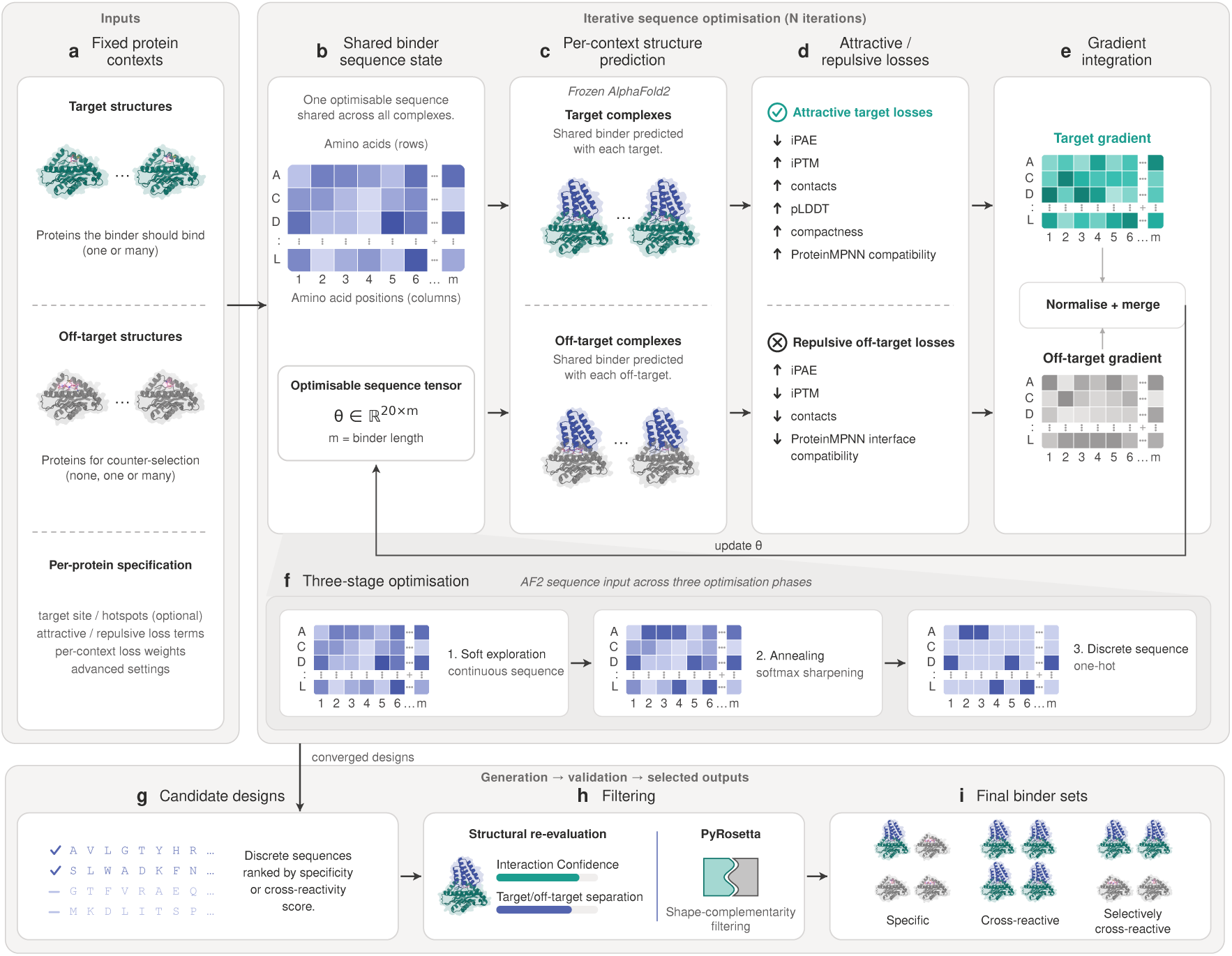
Odin-Multi workflow for multi-context protein binder design. **a**, Odin-Multi receives one or more fixed target structures and, optionally, off-target structures for counter-selection. Each context can be assigned binding sites or hotspots, attractive or repulsive objectives, context-specific weights and additional design settings. **b,** A single optimisable binder sequence, represented by a shared sequence tensor *θ ∈* R^20^*^×m^* for a binder of length *m*, is used across all complexes. **c,** At each iteration, frozen AlphaFold2 models independently predict the shared binder in complex with each target and off-target. **d,** Attractive on-target objectives favour low iPAE, interface contacts, binder confidence and compactness. Repulsive off-target objectives discourage confident and well-packed off-target interfaces. Optional campaign-specific LigandMPNN or ProteinMPNN losses encourage or discourage sequence–structure compatibility. **e,** Context-specific optimisation signals are weighted and combined to update the shared sequence state. **f,** During gradient-based optimisation, the sequence remains parameterised in logit space, while the representation passed to AlphaFold2 progresses through relaxed-logit, temperature-annealed softmax and straight-through one-hot stages. **g,** Converged discrete sequences are ranked according to the desired specificity or cross-reactivity profile. **h,** Candidates selected for experimental testing are re-evaluated with AlphaFold3 and filtered using the interaction prediction score from aligned errors (ipSAE), target-to-off-target iPAE separation and PyRosetta-based shape complementarity. **i,** The workflow yields binder sets with user-defined interaction profiles, including specific, cross-reactive and selectively cross-reactive designs. Teal denotes targets, grey denotes off-targets, and blue denotes the designed binder.

### 2.2 Odin-Multi shifts design populations towards cross-reactivity and specificity

We first evaluated Odin-Multi in a cross-reactivity benchmark across the three class B1 GPCRs, GLP-1R, GCGR and GIPR. We reasoned that since therapeutic development has increasingly progressed from selective receptor agonists towards single peptides with activity across multiple members of this receptor family, including dual GLP-1R/GIPR and triple GLP-1R/GIPR/GCGR agonists, design of a single peptide able to engage multiple receptors would be directly applicable [31, 25, 26]. Notably, the three GPCRs share pairwise sequence identities of only 45.6–51.2% and C*α* RMSDs of 3.49–4.44 Å (Fig. **2**a, left). Despite these differences, joint optimisation via Odin-Multi enriched candidates predicted to bind both receptors in each pair. For the primary AF2 benchmark analysis, each trajectory was evaluated at its final optimisation iteration, rather than selecting the best-performing iteration retrospectively. The same checkpoint was used for the joint- and single-target campaigns; in the single-target campaigns, the second receptor was monitored but did not contribute to sequence optimisation. When requiring both receptor iPTM values to exceed 0.5, 596/616 (96.8%), 999/1,102 (90.7%), and 921/1,103 (83.5%) of designs jointly optimised against GLP-1R–GCGR, GLP-1R–GIPR, and GCGR–GIPR, respectively, passed the threshold. The corresponding single-target pass rates were 18.8% and 36.3% for GLP-1R–GCGR, 10.7% and 18.6% for GLP-1R–GIPR, and 10.6% and 6.8% for GCGR–GIPR. This enrichment persisted across the full range of evaluated iPTM thresholds (Fig. **2**b). Independent AF2 re-evaluation retained enrichment for jointly optimised designs across all three receptor pairs, although its magnitude varied between pairs (Supplementary Fig. 1).

**Figure 2:**
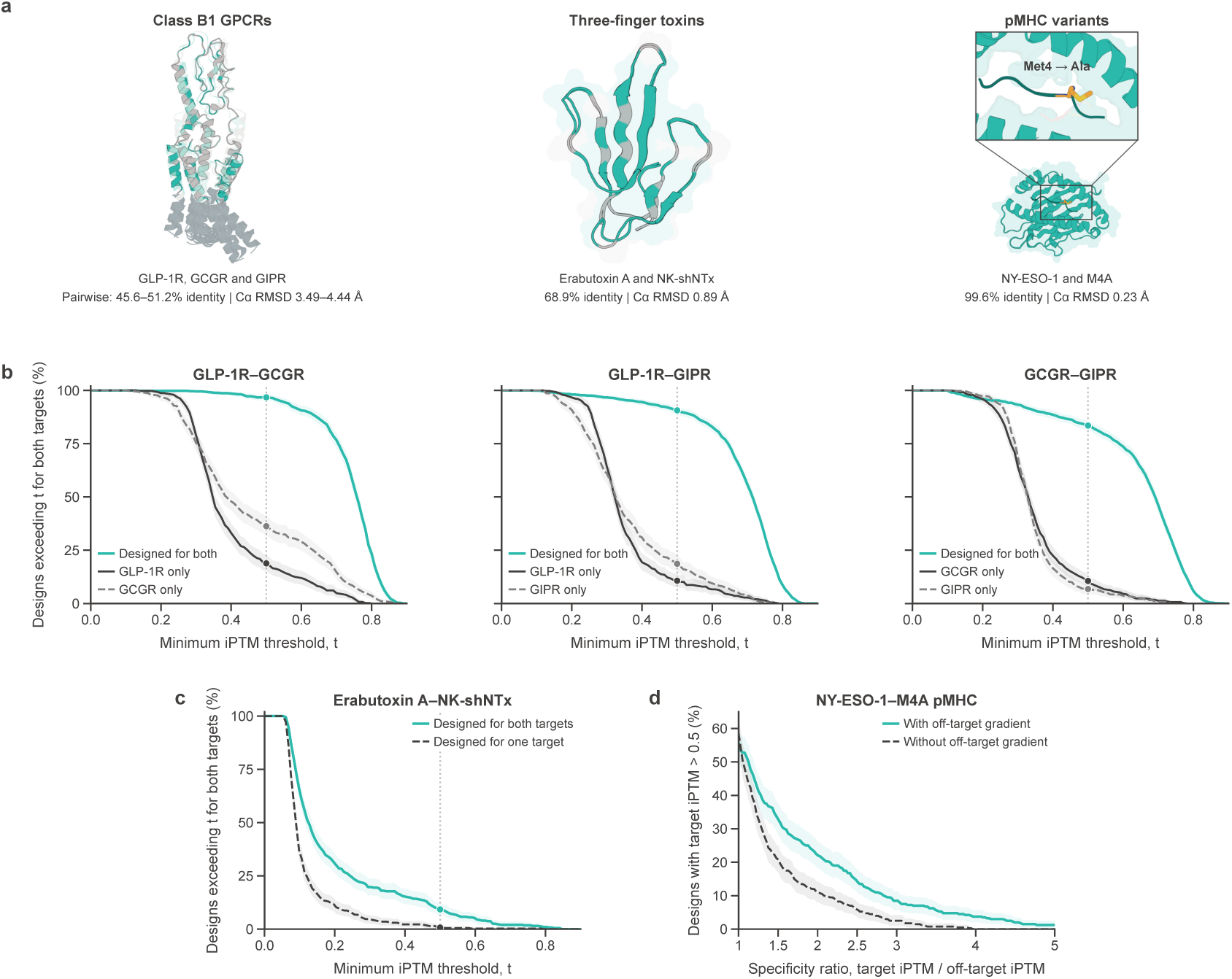
Odin-Multi shifts binder design populations towards cross-reactivity or specificity. **a**, Structural comparison of the three benchmark systems. For the GPCR structures, darker grey indicates receptor regions that were excluded from the cropped structures used for design. Class B1 GPCRs GLP-1R, GCGR and GIPR share 45.6–51.2% pairwise sequence identity and C*α* RMSDs of 3.49–4.44 Å. Erabutoxin A and NK-shNTx share 68.9% sequence identity and a C*α* RMSD of 0.89 Å. The target and off-target pMHC complexes differ by a single peptide residue, Met4*→*Ala, corresponding to 99.6% overall sequence identity and a C*α* RMSD of 0.23 Å. **b,** Percentage of designs for which both receptor iPTM values exceeded the indicated threshold, *t*. For each trajectory, the reported scores were obtained from the final optimisation iteration. No retrospective selection based on either receptor score was performed. The same checkpoint was used for the joint- and single-target campaigns; in the single-target campaigns, the second receptor was monitored but did not contribute to sequence optimisation. The dotted line and markers indicate *t* = 0.5, and the keys report sample sizes and percentages at this threshold. **c,** Percentage of designs exceeding a minimum iPTM threshold for both Erabutoxin A and NK-shNTx. Designs jointly optimised against both toxins retained predicted binding to Erabutoxin A while showing improved predicted binding to the second toxin relative to designs optimised against Erabutoxin A alone. **d,** Percentage of all pMHC minibinder designs satisfying target iPTM *>* 0.5 and the indicated target-to-off-target iPTM ratio threshold. No single ratio threshold was pre-specified. The primary comparison used a global permutation test of the complete yield curves over 160 evenly spaced ratio thresholds from 1 to 5, yielding a maximum absolute between-condition difference of 13.50 percentage points (Holm-adjusted *P* = 0.00070). At a post hoc illustrative ratio cutoff of 2.5, 57/400 (14.2%) designs generated with the off-target objective and 24/400 (6.0%) generated without counter-selection satisfied both criteria. Across **b–d**, teal lines denote optimisation in which two interaction contexts contributed to sequence optimisation: two on-targets in **b** and **c**, or one on-target and one off-target in **d**. Black and grey lines denote the corresponding single-context controls; in **b**, the two control lines correspond to optimisation against each receptor individually. Shaded regions show pointwise 95% bootstrap percentile intervals (2,000 resamples) and describe uncertainty at individual thresholds rather than between-condition significance.

We next tested whether Odin-Multi could promote cross-reactive binding to two short-chain 3FTxs, Erabutoxin A and cobrotoxin-c from *Naja kaouthia* (UniProt ID P59276; hereafter NK-shNTx), which share 68.9% sequence identity and a C*α* RMSD of 0.89 Å (Fig. **2**a, centre). Minibinder sequences were optimised either against Erabutoxin A alone or against both toxins jointly, and predicted interface confidence was evaluated independently for each complex using iPTM. At an iPTM threshold of 0.5, 3/400 (0.8%) of the Erabutoxin A-only designs exceeded the threshold for both targets, compared with 37/400 (9.2%) of the jointly optimised designs (Fig. **2**c). This gain arose from improved predicted binding to the second target while maintaining performance on Erabutoxin A (Supplementary Fig. 2a). A population-level shift was retained following AF3 re-evaluation (Supplementary Fig. 2c), supporting the robustness of the predicted cross-reactivity signal across structure-prediction models. These results demonstrate that joint optimisation enriches designs predicted to bind both 3FTxs, a profile rarely obtained when optimising against Erabutoxin A alone.

Finally, we tested whether an explicit off-target objective could promote discrimination between highly similar pMHC complexes. We optimised against the NY-ESO-1 epitope SLLMWITQC presented on HLA-A*02:01 while counter-selecting against the single-residue variant SLLAWITQC (M4A), which shares the same MHC scaffold and differs by only one peptide residue (99.6% overall sequence identity; C*α* RMSD 0.23 Å; Fig. **2**a, right). Keeping all other settings fixed, the off-target objective shifted the design population towards greater predicted target–off-target separation while broadly preserving on-target confidence (Supplementary Fig. 2b). Because no single specificity threshold was pre-specified, we compared the complete target-qualified specificity-yield curves across target-to-off-target iPTM ratios from 1 to 5 (Fig. **2**d). The curves differed between conditions, with a maximum absolute difference of 13.50 percentage points (Holm-adjusted *P* = 0.00070). At the post hoc illustrative ratio threshold of 2.5, 57/400 (14.2%) counter-selected designs passed, compared with 24/400 (6.0%) designs generated without counter-selection. Independent AF3 re-evaluation was then used to assess whether the separation between conditions was retained under a different structure-prediction model. At the same ratio threshold of 2.5, 59/400 (14.8%) counter-selected designs passed compared with 38/400 (9.5%) designs generated without counter-selection; across the complete yield curves, the maximum absolute difference was 7.50 percentage points (Holm-adjusted global permutation *P* = 0.0612; Supplementary Fig. **2**d).

Together, these computational benchmarks show that Odin-Multi can shift design populations towards either cross-reactivity or on-target/off-target discrimination within a common optimisation framework. Statistical comparisons for the primary AF2 analyses and the independent re-evaluations are reported in Supplementary Table 1.

### 2.3 Joint optimisation yields a dual-target three-finger toxin minibinder

We next applied Odin-Multi in the cross-reactive mode, where a single binder was designed to engage two related targets rather than discriminate between them. We decided to focus on two short neurotoxins, which belong to the 3FTx family and as such are some of the most lethal snake venom toxins. These toxins make ideal design targets, as first 3FTxs, while highly divergent on the sequence level, are sufficiently structurally conserved to merit a polyspecific binder approach and second *de novo* designed neutralisers have so far been designed against individual toxins rather than explicitly optimised for joint recognition of multiple homologues [28]; overcoming this hurdle is key in developing de novo designed next generation antivenoms, as the complexity of snake venoms and multi-toxin nature requires cross-reactive neutralisers in order to remain affordable and manufacturable. Specifically, we proceeded with Erabutoxin A and NK-shNTx from the computational benchmark, since they shared only 68.9% sequence identity, providing a non-trivial cross-reactive design task (Fig. **3**a). Odin-Multi was run in cross-reactive mode, with both toxins treated as on-targets and assigned equal context weights, thereby allowing both predicted complexes to contribute directly to optimisation of the shared minibinder sequence.

**Figure 3:**
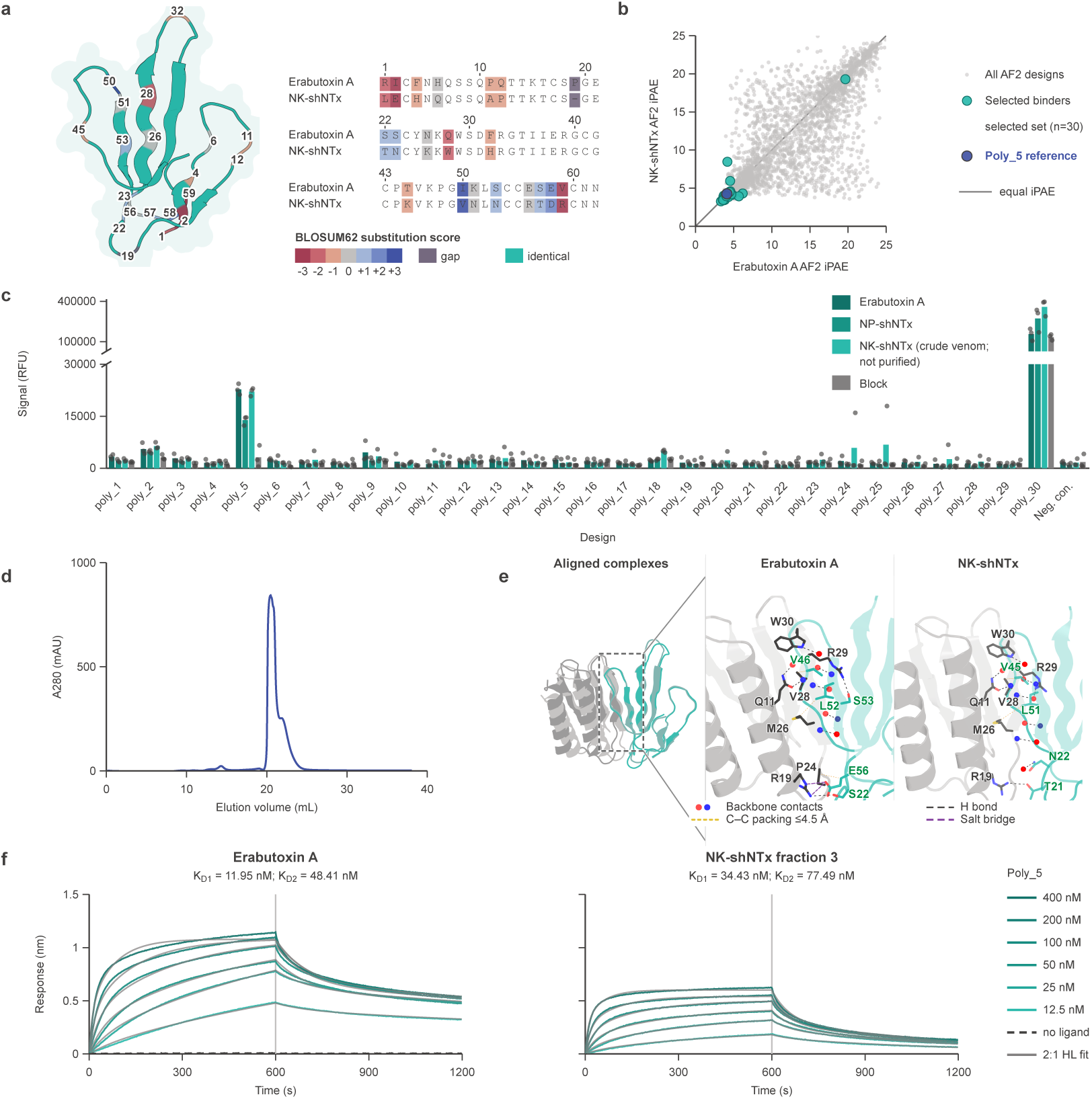
Experimental validation of an Odin-Multi-designed dual-target 3FTx minibinder. **a**, Structural and sequence comparison of Erabutoxin A and NK-shNTx. Residue substitutions are coloured according to their BLOSUM62 scores. **b,** AF2-derived iPAE values for designs evaluated independently against Erabutoxin A and NK-shNTx. Grey points represent all generated designs, whereas teal points represent the 30 designs selected for experimental testing. A single design, Poly_5 (highlighted in purple), reached elevated binding signals across the tested toxins (panel **c**) and was selected for further investigation. The diagonal line denotes equal AF2 iPAE for the two targets. Although AF2 metrics are shown here, final designs were selected following AF3 re-evaluation, requiring *ipSAE >* 0.61 and Rosetta interface shape complementarity greater than 0.5 for both targets. **c,** DELFIA screening of the purified minibinders against Erabutoxin A, purified *Naja pallida* short neurotoxin 1 (NP-shNTx), and *Naja kaouthia* whole venom, together with a skimmed-milk blocking control. NP-shNTx was included as the closest available purified homologue of NK-shNTx because purified NK-shNTx was unavailable. Bars show mean fluorescence signals and grey points indicate individual replicates. Poly_5 showed elevated signals against the toxin-containing coatings relative to the blocking control, whereas Poly_30 produced high signals against all coatings, including the blocking control, indicating non-specific reactivity. The discontinuous y-axis accommodates the substantially higher signals observed for Poly_30. One anomalously high replicate for Poly_13 against NP-shNTx was excluded from the analysis. **d,** Size-exclusion chromatography profile of Poly_5 following C-tag affinity purification. **e,** AF2-predicted Poly_5 complexes with Erabutoxin A and NK-shNTx, shown following alignment of the toxin backbones, together with close-up views of the predicted interfaces. Selected interface residues are shown as sticks and labelled. Red and blue dots mark predicted backbone contacts, and dashed lines denote predicted hydrogen bonds, carbon–carbon packing within 4.5 Å, and salt bridges, as indicated in the panel. **f,** Biolayer-interferometry sensorgrams for Poly_5 binding to Erabutoxin A and candidate NK-shNTx-containing fraction 3 across a concentration series from 12.5 to 400 nM. Data were fitted using a 2:1 heterogeneous-ligand model with independent Rmax, yielding *K*_D1_ = 11.95 nM and *K*_D2_ = 48.41 nM for Erabutoxin A and *K*_D1_ = 34.43 nM and *K*_D2_ = 77.49 nM for fraction 3.

Thirty designs that passed AlphaFold3 filters were cloned, expressed, and purified (Fig. **3**b). Twenty-four of the 30 designs produced protein yields above that of the empty-plasmid negative control (Supplementary Fig. **3**a). The purified minibinders were evaluated by DELFIA against Erabutoxin A, purified *Naja pallida* short neurotoxin 1 (NP-shNTx), and *Naja kaouthia* whole venom, together with a skimmed-milk blocking control. Because purified NK-shNTx had become commercially unavailable, NP-shNTx was included as the closest available purified homologue of the designed *N. kaouthia* target, sharing 85.2% sequence identity. One design, Poly 5, showed elevated binding signals across all three toxin-containing coatings, including signal-to-block ratios exceeding fourfold for Erabutoxin A and *N. kaouthia* whole venom (Fig. **3**c). This corresponded to an experimental hit rate of 1/30 designs tested (3.3%) or 1/24 among designs that expressed above the empty-plasmid negative-control yield (4.2%). Although the screening conditions and intended interaction profiles differed, this hit rate was comparable to the 1/44 design hit rate we reported for single-target short-chain neurotoxin minibinders previously [28]. One further design, Poly 30, bound all coatings, including the blocking control, indicating non-specific reactivity rather than family-level recognition, and was excluded from further analysis. The remaining designs showed no or limited target-associated signal relative to background.

RP-HPLC fractionation and MALDI-TOF analysis identified fraction 3 as the candidate NK-shNTx-containing fraction used for subsequent binding analysis (Supplementary Figs. **3**b–c and **4**; Methods). Prior to BLI, Poly 5 was purified by C-tag affinity purification and size exclusion chromatography (SEC), yielding a single peak with a modest shoulder (Fig. **3**d), while SDS-PAGE showed a predominant band at the expected molecular mass (Supplementary Fig. **3**d), consistent with a predominantly monomeric species. Binding of Poly 5 to Erabutoxin A and the candidate NK-shNTx-containing fraction 3 was measured using biolayer interferometry (BLI) (Fig. **3**f). The higher-affinity binding component of a 2:1 heterogeneous-ligand model yielded apparent *K*_D1_ values of 11.95 nM for Erabutoxin A and 34.43 nM for fraction 3 (Fig. **3**f; Supplementary Table 2).

**Figure 4:**
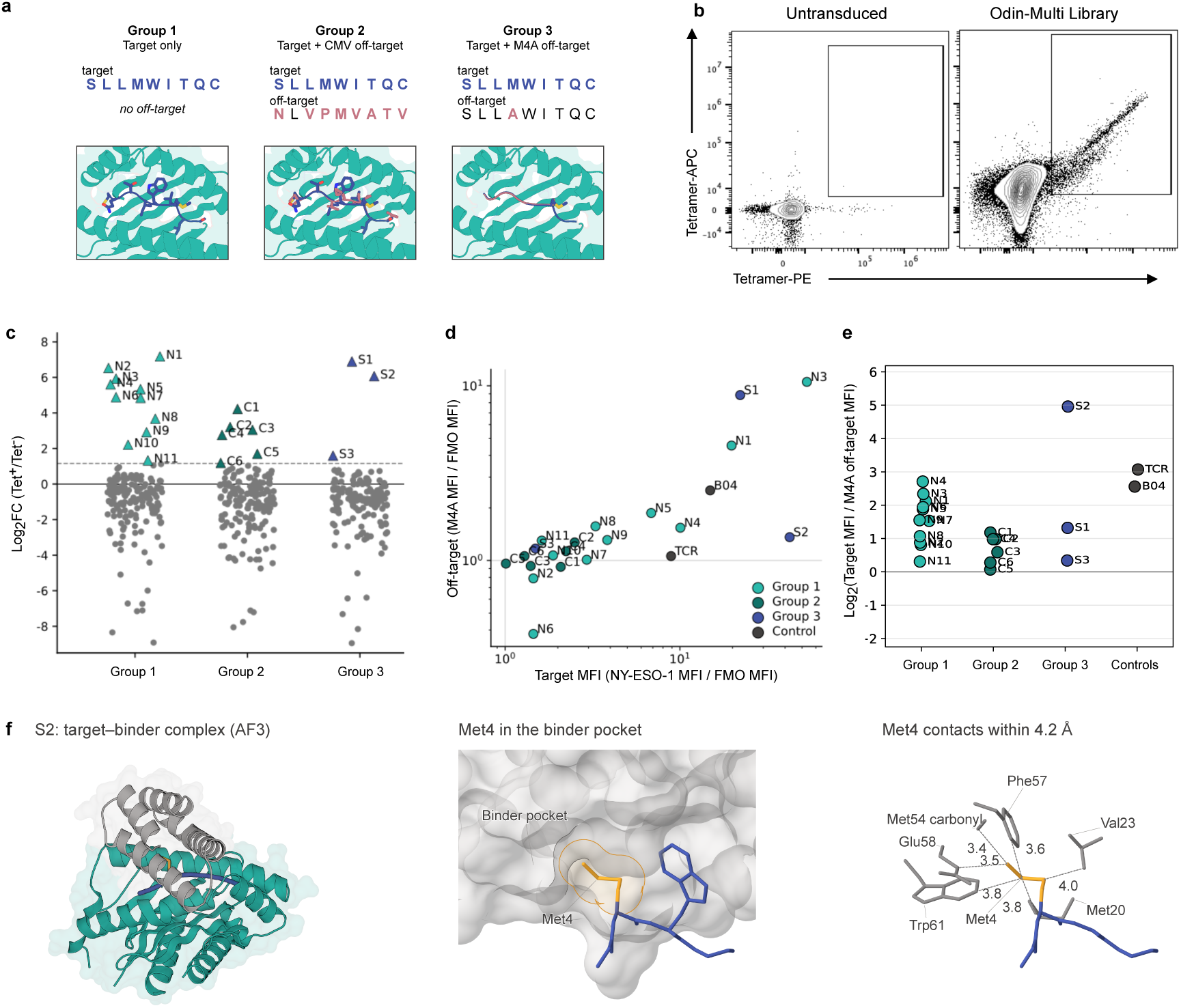
Experimental validation of pMHC targeting miBds designed using Odin-Multi. **a**, Schematic of the three Odin-Multi design conditions used for the pMHC specificity campaign. Group 1 was optimised against the NY-ESO-1 target, SLLMWITQC/HLA-A*02:01, without an explicit off-target. Group 2 was optimised against the same target while penalising binding to empty HLA-A*02:01 and the sequence-divergent CMV pMHC, NLVPMVATV/HLA-A*02:01. Group 3 was optimised against the target while penalising binding to empty HLA-A*02:01 and the near-identical M4A variant, SLLAWITQC/HLA-A*02:01. **b,** The pooled library contained 166 designs from the target-only condition and 167 designs from each of the CMV- and M4A-counter-selected conditions, for a total of 500 miBds. The Odin-Multi-designed miBds were cloned into a 41BB–CD3*ζ* CAR vector and lentivirally transduced into CD3 KO Jurkat cells, enabling mammalian cell-surface display. The library was stained with SLLMWITQC/HLA-A*02:01 tetramers labelled with PE and APC. Flow plots show the sorted cells gated on singlet/live/tNGFR^+^ cells. **c,** The top 20 designs were selected for single-clone characterisation, with the lowest-ranked selected design having a Log_2_FC of 1.2 (triangles). These comprised 11 designs from the target-only condition (Group 1, N1–11), six from the CMV-counter-selected condition (Group 2, C1–6), and three from the SLLAWITQC (M4A)-counter-selected condition (Group 3, S1–3). The dotted line marks Log_2_FC of 1. **d,** Dot plot showing individual tetramer staining of the 20 selected clones. Clones were stained with the NY-ESO-1 target tetramer (SLLMWITQC/HLA-A*02:01) and the M4A off-target tetramer (SLLAWITQC/HLA-A*02:01) separately. MFI values were normalised to the tetramer FMO control, with values above 1 (grey lines) indicating signal above background. **e,** Relative antigen discrimination of the screened clones. Discrimination was quantified as the log_2_ ratio of NY-ESO-1 target MFI to M4A off-target MFI. Values above zero indicate preferential recognition of the target antigen, while values close to zero indicate similar binding to the target and off-target variant. The grey line marks equal target and off-target binding. **f,** AlphaFold3-predicted structure of S2 bound to SLLMWITQC/HLA-A*02:01. Left, overview of the complex, with the minibinder shown in grey, HLA in teal, and the presented peptide in blue. Middle, closeup showing the peptide Met4 side chain, highlighted in orange, accommodated within a pocket on the minibinder surface. Right, predicted contacts within 4.2 Å between Met4 and the minibinder, involving Met20, Val23, the Met54 backbone carbonyl, Phe57, Glu58, and Trp61. Dashed lines indicate selected interatomic distances, labelled in Å. Residue numbering refers to the peptide for Met4 and to the minibinder for the surrounding residues.

Structural analysis of the predicted Poly 5 complexes suggested that the minibinder engages the exposed *β*-sheet face of both toxins through a largely conserved, backbone-dominated interaction pattern (Fig. **3**e). In both models, the corresponding toxin residues V46/V45, I50/V49, L52/L51, and C54/C53 formed similar predicted hydrogen-bonding and packing contacts with Poly 5, where residues are reported as Erabutoxin A/NK-shNTx. Beyond this shared core, each model contained a few additional contacts, each involving different regions of the toxin: S53 and E56 in Erabutoxin A, and T21 and N22 in NK-shNTx. The predicted binding mode was therefore the same for both toxins, with the sequence differences between them accommodated by contacts outside the shared *β*-sheet face.

These results demonstrate that Odin-Multi supports targeted cross-reactivity as a design objective and yielded a minibinder able to bind both to Erabutoxin A and a candidate NK-shNTx-containing venom fraction with high affinity.

### 2.4 Counter-selection produces a class I pMHC minibinder with greater antigen discrimination

We next sought to experimentally validate Odin-Multi in the specificity regime, using a stringent discrimination problem with an established experimental benchmark. Selective recognition of class I MHC-presented peptides is critical given the risk of targeting healthy tissue, as highlighted when TCRs targeting MAGE-A3-derived peptides showed fatal cross-reactivity towards peptides derived from the cardiac protein TITIN [4]. Because pMHC off-targets often share the same MHC scaffold and differ from the on-target only in the presented peptide [18, 16], discrimination must distinguish subtle differences in the peptide-exposed surface rather than global structural differences.

We focused on the NY-ESO-1 epitope, SLLMWITQC, presented on HLA-A*02:01, for which an experimentally validated *de novo* minibinder, NY1-B04, has been reported [18]. NY1-B04 binds the single-residue variant SLLAWITQC/HLA-A*02:01 (M4A) at approximately wild-type-equivalent avidity, providing both a demanding specificity target and a direct point of comparison for designs generated under an explicit off-target objective. We designed minibinders under three conditions of increasing constraint (Fig. **4**a). The first applied no off-target penalty and served as the target-only condition. The second penalised binding to empty HLA-A*02:01 and to a pMHC presenting the sequence-divergent CMV epitope, NLVPMVATV/HLA-A*02:01. The third penalised binding to empty HLA-A*02:01 and to the M4A variant, which differs from the target by a conservative methionine-to-alanine substitution at position four and represents the most stringent of the three tests.

As specificity constraints increased, progressively more design trajectories were required to obtain candidates passing the downstream AF3 filters, reflecting the increasing difficulty of simultaneously satisfying attractive target and repulsive off-target objectives. We ran 203 design trajectories for the target-only condition, 453 for the CMV-counter-selected condition, and 1,700 for the M4A-counter-selected condition, of which 155, 341, and 1,235, respectively, completed without early termination and yielded designs. Early-termination rates were comparable across the three conditions (24%, 25% and 27%), indicating that the additional trajectories reflect lower downstream AF3 filter pass rates rather than a higher rate of trajectory failure.

A total of 166 designs from the target-only condition and 167 designs from each of the CMV- and M4A-counter-selected conditions were tested experimentally using a mammalian surface-display system in which minibinders (miBds) were expressed on TCR-deficient Jurkat T cells. The three design conditions were screened together as a pooled library of 500 miBds. The library-transduced cells were stained with pMHC tetramers, FACS sorted as tetramer-positive and tetramer-negative populations (Fig. **4**b). Based on log_2_ FC enrichment values (Fig. **4**c), the top 20 designs were selected for single-clone characterisation, with the lowest-ranked selected design having a log_2_ FC of 1.2. These comprised 11/166 designs from the target-only condition, 6/167 from the CMV-counter-selected condition, and 3/167 from the M4A-counter-selected condition. These twenty designs were evaluated individually for SLLMWITQC/HLA-A*02:01 binding avidity by pMHC tetramer staining and in parallel evaluated for binding to the off-target peptides included in the design campaign.

Staining of the twenty individual clones with the SLLMWITQC/HLA-A*02:01 tetramer confirmed that Odin-Multi generated target-binding clones under all three design conditions (Fig. **4**d, Supplementary Fig. **5** a). The target-only condition (Group 1) produced the highest number of target-binding minibinders, with all 11 selected clones showing some degree of binding, including three (N1, N3, N4) with higher MFI than the SLLMWITQC/HLA-A*02:01-targeting TCR clone 1G4 (Supplementary Fig. **5**c). Several of these clones also recognised the M4A mutant, most notably N1 and N3, the two highest-avidity target binders in the group, which bound the off-target with comparably high avidity. The CMV-counter-selected condition (Group 2) primarily produced low-avidity target binders, whereas all Group 3 clones retained detectable target binding, with S1 and S2 exceeding both clone 1G4 and NY1-B04 in MFI (Supplementary Fig. **5**c).

CMV recognition was detected for a single target-only clone, N8, and for none of the counter-selected clones in either group (Supplementary Fig. **5**b–c). CMV cross-reactivity was therefore possible but uncommon in the absence of explicit counter-selection, leaving limited dynamic range in which to quantify a benefit from the CMV objective.

Antigen discrimination was quantified as the log_2_ ratio of NY-ESO-1 target MFI to M4A off-target MFI, where positive values indicate preferential recognition of the target (Fig. **4**e). Consistent with the avidity data, the strongest target binders from the target-only condition were among the weakest discriminators, and clones from the target-only and CMV-counter-selected conditions showed moderate to limited target/M4A separation overall. The M4A-counter-selected condition produced one clone, S2, that both exceeded NY1-B04 in target avidity and discriminated more effectively between the two pMHCs, which share 99.6% sequence identity and differ by a single conservative substitution at peptide position four [18]. S2 also exceeded the TCR control, although this likely reflects the lower target avidity of the TCR rather than reduced TCR specificity. The remaining M4A-counter-selected clones, S1 and S3, showed only modest discrimination.

To explore a possible structural basis for S2 discrimination, we examined its AF3-predicted complex with the NY-ESO-1 target pMHC (Fig. **4**f). The peptide Met4 side chain occupies a pocket on the minibinder surface, with predicted contacts within 4.2 Å involving Met20, Val23, the Met54 backbone carbonyl, Phe57, Glu58, and Trp61. This arrangement suggests that replacing methionine with the smaller alanine side chain could reduce packing interactions within the pocket, providing a possible explanation for the lower M4A tetramer staining observed for S2. This interpretation is based on the predicted target-bound structure and does not establish the binding geometry of the M4A complex.

Counter-selection carried a yield cost: designs exceeding the enrichment threshold fell from 11/166 in the target-only condition to 6/167 and 3/167 in the CMV- and M4A-counter-selected conditions, and Groups 2 and 3 differed in both off-target identity and off-target weight, so their comparison does not isolate the contribution of either factor. From the smallest of these sets, however, came S2: a design that stained at higher MFI than the published NY1-B04 prototype while also discriminating markedly better between the target and the single-residue M4A variant [18]. Distinguishing two surfaces this similar is among the hardest discrimination tasks in current binder design, and here it was obtained by specifying the off-target as a design objective rather than relying solely on downstream screening for it.

## 3 Discussion

Odin-Multi encodes multi-target and multi-off-target interaction profiles directly into the optimisation objective rather than as a downstream filter. Multistate protein design has long treated specificity as simultaneous positive and negative design across desired and undesired structural states [21, 22, 23], generally using discrete sequence search under physics-based energy functions. More recently, AlphaDesign extended AlphaFold-based *de novo* design to multistate and multitarget objectives by evolving discrete amino-acid sequences under combined AlphaFold-derived fitness functions [24]. Odin-Multi instead builds on the continuous AlphaFold-based hallucination framework used by BindCraft [15], extending its single-complex optimisation to a shared sequence representation evaluated across multiple independently predicted complexes. Context-specific gradients are combined during optimisation, allowing attractive objectives for on-targets and repulsive objectives for specified off-targets to shape the same sequence. Because these contexts contribute during sequence optimisation, specificity or cross-reactivity can alter which sequences are generated, rather than being used only to reject undesired candidates after generation. The effect was visible at the population level: only 0.8% of minibinders optimised against Erabutoxin A alone exceeded the dual-target iPTM threshold compared with 9.2% under joint optimisation, while at an illustrative target-to-off-target iPTM ratio threshold of 2.5, pMHC specificity yield increased from 6.0% without counter-selection to 14.2% with counter-selection. These shifts are important because downstream selection can only act on interaction profiles already represented among the generated candidates.

In the cross-reactive mode, Odin-Multi produced Poly 5, a minibinder that bound both a purified short-chain neurotoxin, Erabutoxin A, and a candidate NK-shNTx-containing fraction of *Naja kaouthia* venom with apparent nanomolar affinity. Structural prediction suggests how a single sequence accommodates both: the minibinder engages the same exposed *β*-sheet face in each complex through a largely backbone-mediated core, with sequence differences between the toxins accommodated by peripheral contacts. This resembles the conserved-epitope recognition sought in broadly neutralising antitoxin antibodies and nanobodies [20, 27], rather than independent recognition of two unrelated surfaces. If this recognition mode generalises to further family members, and if binding translates into neutralisation, it could provide a route towards family-covering binders for next-generation antivenoms [5, 28].

The pMHC results illustrate this distinction in a particularly stringent setting. Recent *de novo* pMHC binder studies have addressed specificity predominantly after candidate generation: through downstream cross-panning [18], ProteinMPNN- and AF2-based assessment against identified off-target peptides [16], or, for an *α*-helical TCR mimic against the NY-ESO-1 C9V/HLA-A*02 system, by searching naturally presented HLA-A*02 peptides for potential off-targets followed by structural prioritisation and experimental testing [29]. Odin-Multi instead incorporates a defined off-target directly into sequence optimisation. The lead design S2 distinguished the NY-ESO-1 on-target from the single-residue M4A off-target more effectively than the previously reported *de novo* minibinder NY1-B04 under the same tetramer-staining assay [18]. This provides experimental evidence that explicitly penalising predicted binding to a closely related off-target during optimisation can improve antigen discrimination.

The pMHC campaigns also highlight practical considerations for explicit multi-context optimisation. Computational cost scales approximately additively with the number of interaction contexts, because each target or off-target is predicted and evaluated independently during optimisation. Consistent with this, increasing the pMHC objective from one context in the target-only condition to three contexts in the counter-selected conditions increased median per-trajectory runtime approximately threefold on matched V100 hardware, from 20.2 min to 59.3–61.5 min. Multi-context optimisation can impose a second cost through reduced design yield, because candidates must satisfy the desired interaction criteria across multiple contexts simultaneously. Accordingly, the M4A campaign required 1,700 trajectories to assemble its screened design set compared with 203 trajectories for the target-only campaign, reflecting the lower downstream AF3 filter pass rate under the additional specificity constraints. Beyond computational cost, counter-selection is most informative when the baseline designs show measurable binding to the competing off-target. In the pMHC experiments, CMV recognition appeared in a single target-only clone, N8, and in none of the CMV-counter-selected clones, leaving little dynamic range in which an additional benefit could have been detected. Choosing an off-target that represents a plausible competing interaction among otherwise viable target-binding designs therefore matters as much as specifying an off-target at all.

Several limitations qualify these conclusions. Each experimentally tested design regime yielded a single lead with the intended interaction profile, and broader testing will be required to establish how consistently the approach improves experimental success; success rates vary widely across targets and campaigns in current *de novo* binder-design pipelines [15, 28]. In the 3FTx campaign, the RP-HPLC behaviour and intact mass of fraction 3 were consistent with the short-chain three-finger neurotoxins identified in previous *N. kaouthia* venomics analyses [32], although the observed mass matched neither candidate exactly; intact-mass MALDI-TOF cannot establish sequence identity, and binding therefore cannot be assigned definitively to NK-shNTx. Computational evaluation was also sensitive to the prediction setup. AF2 has been reported to be relatively insensitive to point mutations at protein–protein interfaces, and AF3-based filtering can itself retain false-positive interaction predictions [15], making the M4A benchmark, in which target and off-target differ by a single peptide residue, a demanding use case for structure-prediction-based optimisation. The specificity separation observed with AF2 was attenuated on AF3 re-evaluation (Holm-adjusted *P* = 0.0612), underscoring the model dependence of the predicted effect. We therefore interpret the computational separation as supporting rather than definitive evidence that counter-selection altered the design population, with the discrimination observed for S2 providing orthogonal experimental support, albeit from a single lead. The GPCR cross-reactivity signal was likewise retained under an independent AF2 setup using AF2-Monomer-predicted receptor structures, although its magnitude differed between prediction setups. This sensitivity is consistent with known limitations in AF2 modelling of GPCR conformational states, including biases related to the conformational distribution represented in the training data [33].

Beyond the specific systems addressed here, Odin-Multi applies to binder-design problems in which the desired output is an interaction profile rather than a single interaction. Multi-specific antibody engineering, where simultaneous recognition of several antigens is required, maps naturally onto the cross-reactive regime [2, 3]; receptor subtype discrimination, where closely related receptors share conserved binding surfaces, maps onto the exclusion regime; and family-targeting reagents, where broad recognition of related proteins is desired while unrelated cross-reactivity is not, may require both modes simultaneously. As *de novo* binder design advances towards clinical development and practical application, methods that directly control patterns of target and off-target interactions during generation should complement continued improvements in affinity, developability, prediction accuracy and downstream specificity assessment.

## 4 Methods

### 4.1 Odin-Multi computational design

#### 4.1.1 On-target and off-target loss functions

Each predicted binder–target complex was scored using AF2-derived confidence metrics and geometric constraints [11, 12]. For on-targets, the objective included iPAE (weight 0.1), an interface-contact loss (weight 1.0; 20 Å cutoff; minimum of two contacts), intra-binder PAE (weight 0.4), an intra-binder contact loss (weight 1.0; 14 Å cutoff), binder pLDDT (weight 0.1), radius of gyration (weight 0.3), helicity (weight *−*0.3), and pTMEnergy (weight 0.05) [34]. Off-target loss formulations were campaign specific and are described in the corresponding benchmark sections below.

#### 4.1.2 Sequence–structure regularisation

Sequence–structure regularisation was campaign-specific. The LigandMPNN- and ProteinMPNN-derived regularisation terms used across campaigns are defined below, while their campaign-specific use and weights are summarised in Table 1. Conventional post hoc MPNN redesign was disabled throughout.

**Table 1:** Campaign-specific design and evaluation settings used in Odin-Multi.

| *Computational benchmarks* |  |  |  |
| --- | --- | --- | --- |
|  | GPCR | 3FTx | pMHC |
| **Context weights** | Joint: equal across both receptors.<br>Single-target: remaining receptors monitored only, no gradient contribution. | Joint: equal across both toxins.<br>Single-target: Erabutoxin A only. | Counter-selected: M4A off-target at 0.30, repulsive.<br>Target-only: no off-target context. |
| **Optimisation stages** | 75 logits-to-soft;<br>40 temperature-annealed softmax;<br>25 straight-through one-hot;<br>10 PSSM-guided semigreedly. | 75 logits-to-soft;<br>45 temperature-annealed softmax;<br>20 straight-through one-hot. | 75 logits-to-soft;<br>45 temperature-annealed softmax;<br>5 straight-through one-hot, evaluating and differentiating through all five AF2-Multimer models at each iteration rather than one sampled model. |
| **Regularisation** | LigandMPNN (v_32_010) sequence-distribution loss at 1.0, evaluated independently per on-target. | None. | None. |
| **Independent re-evaluation** | AF2-Multimer, with AF2-Monomer receptor structures as initial guess. Applied to the minimum-loss retained sequence rather than the final iteration. | AF3. | AF3. |
| **Designs analysed** | Joint: 616 GLP-1R-GCGR; 1,102 GLP-1R-GIPR; 1,103 GCGR-GIPR.<br>Single-target: 393 GLP-1R; 463 GCGR; 381 GIPR. | Joint: 400.<br>Single-target: 400. | Counter-selected: 400.<br>Target-only: 400.<br>Both capped chronologically by completion time. |
| *Experimental design campaigns* |  |  |  |
|  | 3FTx | pMHC |  |
| **Context weights** | Joint: equal across both toxins. | Target-only: no off-target context.<br>CMV: CMV pMHC at 0.20, empty HLA-A*02:01 at 0.01, both repulsive.<br>M4A: M4A pMHC at 0.30, empty HLA-A*02:01 at 0.01, both repulsive. |  |
| **Optimisation stages** | 75 logits-to-soft;<br>45 temperature-annealed softmax;<br>20 straight-through one-hot. | 75 logits-to-soft;<br>45 temperature-annealed softmax;<br>20 straight-through one-hot. |  |
| **Regularisation** | On-target: ProteinMPNN structural-preference loss at 0.1; sequence-distribution loss at 0.05.<br>Both evaluated independently per on-target. | On-target: ProteinMPNN structural-preference loss at 0.1; sequence-distribution loss at 0.05.<br>Off-target: structural-preference loss at $-0.1$, soft distance-based interface weights; sequence-distribution loss at $-0.05$, hard interface mask. | |
| **Candidate selection and filtering** | Three best AF2 candidates from the final 20 iterations per trajectory, ranked by $S_{\text{cross}}$. Required for both toxins: $ipSAE > 0.61$; shape complementarity $> 0.5$. | Three best AF2 candidates from the final 20 iterations per trajectory, ranked by $S_{\text{spec}}$. Required: off-target-to-target iPAE ratio $> 1.5$; $ipSAE > 0.45$, shape complementarity $> 0.5$.<br>Ranked by the AF3 clipped-iPAE specificity score, at most two sequences per trajectory. | |
| **Designs analysed** | Unconstrained: 522 run, 418 yielded scored designs.<br>Hotspot-constrained: 2,000 run, 1,664 yielded scored designs.<br>Screened panel: 30 designs. | Target-only: 203 trajectories run, 155 yielded designs.<br>CMV: 453 run, 341 yielded designs.<br>M4A: 1,700 run, 1,235 yielded designs.<br>Screened library: 166, 167 and 167 designs respectively, 500 total. |  |

For the LigandMPNN sequence-distribution loss, we adapted the inverse-folding sequence-recovery regularisation implemented in Mosaic [35], replacing its ProteinMPNN-based procedure with autoregressive LigandMPNN sampling [30]. A frozen autoregressive LigandMPNN model (v 32 010) was conditioned on the current AF2-predicted target–peptide complex. The target sequence was fixed and placed before the peptide in the decoding order. Sixteen peptide sequences were sampled at a temperature of 0.1, and their one-hot encodings were averaged position-wise to obtain an empirical amino-acid teacher distribution. Categorical cross-entropy between this teacher distribution and the current peptide sequence distribution was added to the objective. During back-propagation, both the sampled teacher distribution and the structural inputs were treated as fixed. Consequently, this loss updated only the peptide sequence logits and did not propagate gradients through LigandMPNN or AF2. For joint-target campaigns, the loss was evaluated independently for each on-target and the resulting terms were combined.

Two ProteinMPNN-derived regularisation terms adapted from ColabDesign and Frank *et al.* were used [36, 14]. The structural-preference loss penalised minibinder positions at which the Protein-MPNN amino-acid distribution lacked a strong preference, quantified as the negative logarithm of the highest amino-acid probability at each position. For on-targets, this loss was evaluated across all minibinder positions. Gradients propagated through the frozen ProteinMPNN model and AF2 to the design parameters, favouring backbones with strong and well-defined ProteinMPNN sequence preferences. For pMHC off-targets, the loss was restricted using soft distance-based interface weights and applied repulsively.

The sequence-distribution loss adapted the corresponding sequence-matching objective as a Kullback– Leibler divergence from the current minibinder sequence distribution to the ProteinMPNN-predicted distribution. During backpropagation, the ProteinMPNN-predicted distribution was treated as a fixed target. Consequently, this loss updated only the minibinder sequence logits and did not propagate gradients through ProteinMPNN or AF2. For on-targets, the loss was evaluated across all minibinder positions. For pMHC off-targets, it was restricted to predicted interface residues using a hard interface mask and applied repulsively, discouraging agreement with the ProteinMPNN-preferred sequence at the off-target interface.

#### 4.1.3 Gradient normalisation and combination

Gradient combination was campaign-specific. In the GPCR computational benchmark, gradients from the two on-target contexts were combined by weighted summation. PCGrad [37] was used in campaigns containing explicit off-target objectives, including the pMHC specificity benchmark and experimental pMHC counter-selection campaigns. The experimental 3FTx design campaign also used PCGrad to combine the two on-target contexts. Single-context target-only campaigns required no cross-context gradient combination. RMS normalisation and context weighting were applied according to the implementation used for each campaign.

Campaign-specific target and off-target context weights are summarised in Table 1.

#### 4.1.4 Sequence optimisation

Sequence optimisation used a three-stage protocol adapted from ColabDesign and BindCraft [36, 15]. First, continuous sequence logits were optimised while the representation supplied to AF2 was progressively transformed towards softmax amino-acid probabilities. This was followed by temperature-annealed softmax optimisation. In the third stage, AF2 received an exact one-hot sequence in the forward pass, while gradients were propagated through the corresponding soft amino-acid distribution using the straight-through estimator. Campaign-specific iteration counts and any subsequent discrete refinement steps are summarised in Table 1.

#### 4.1.5 Candidate ranking

Candidate sequences were collected from the campaign-specific iterations using discrete one-hot sequences in the AF2 forward pass. Unless otherwise specified, cross-reactivity candidates were ranked by their worst-performing on-target:

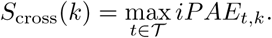

The candidate with the lowest *S*_cross_ is selected, favouring balanced binding across all on-targets. For specificity designs, candidates are ranked by the separation between the worst on-target interaction and the strongest predicted off-target interaction:

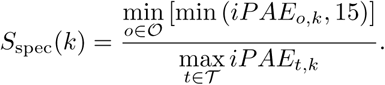

Higher values indicate stronger predicted binding across the on-targets and weaker binding to all off-targets. Off-target *iPAE* values were capped at 15 Å as a pragmatic ranking threshold. Values at or above this level were treated as sufficiently inconsistent with a confident interaction that further increases towards the AF2 PAE ceiling should not provide an additional ranking advantage. Candidates were ranked by *S*_cross_ or *S*_spec_ as defined above. Campaign-specific candidate-collection and downstream selection settings are summarised in Table 1.

#### 4.1.6 Filtering

For the experimental design campaigns, selected designs were re-evaluated using AlphaFold3 (AF3) as separate binary complexes with each on-target and off-target. Where several candidates from the same trajectory were re-evaluated, the candidate minimising the worst-target AF3 iPAE was retained, and only that candidate was carried into downstream filtering. Consequently, the sequence selected for experimental testing was not necessarily the candidate that minimised the corresponding AF2 objective.

Filtering considered target–off-target iPAE separation for specificity campaigns, the interaction prediction score from aligned errors (ipSAE) [38], and PyRosetta-derived interface shape complementarity (SC) [39]. For cross-reactivity campaigns, the on-target filtering criteria were required to be satisfied for every on-target. The ipSAE and shape-complementarity criteria were based on empirical thresholds associated with successful binders reported by Overath *et al.* [40]. Campaign-specific final filtering thresholds are summarised in Table 1.

Designs passing the AF3 filters were subsequently ranked using the campaign-specific selection procedures summarised in Table 1.

#### 4.1.7 Software implementation and computational settings

Odin-Multi is implemented in JAX, extending ColabDesign [36] and BindCraft [15] to support parallel multi-target and off-target optimisation. AF2-Multimer parameters are used with a single recycle during design [11, 12]; at each iteration, one of the five AF2 models is randomly sampled. Computations were run on Azure cloud instances using NVIDIA T4 and V100 GPUs, with one design trajectory per GPU instance. Amino acid biasing, sequence initialisation, and other practical details follow the BindCraft implementation. For the experimental 3FTx and pMHC design campaigns, target and off-target structures were modelled using AlphaFold3 [41] and prepared following the same target-preparation protocol as in BindCraft [15]. The structures and prediction setup used for the GPCR benchmark are described separately below. Campaign-specific design and evaluation settings are summarised in Table 1.

### 4.2 Computational benchmark setup

All computational benchmarks used the common loss, gradient-combination and sequence-optimisation procedures defined in the *Odin-Multi computational design* subsection. Benchmark-specific comparison conditions and datasets are described in the corresponding benchmark subsections, and statistical procedures are described under *Statistical analysis*.

#### 4.2.1 Checkpoint selection and independent re-evaluation

For the primary AF2 benchmark analyses, the GPCR, 3FTx and pMHC trajectories were evaluated at their final optimisation iteration. The same rule was applied to all optimisation conditions within each benchmark, without retrospective selection based on target or off-target prediction scores.

For independent re-evaluation, the final sequences from the 3FTx and pMHC benchmarks were modelled using AlphaFold3. The GPCR AF2 re-evaluation instead used the minimum-loss sequence retained from each trajectory. Consequently, the primary GPCR analysis and independent reevaluation used different sequence-selection rules and could evaluate different sequences from the same trajectory.

These benchmark analyses were separate from the candidate-ranking and filtering procedures used to select designs for experimental testing, which are described in the corresponding experimental-design sections.

#### 4.2.2 GPCR cross-reactivity benchmark

The GPCR benchmark used 20–40-residue peptides, with peptide length randomly sampled for each design trajectory, designed against cropped predicted structures of GLP-1R, GCGR and GIPR. The GLP-1R input contained residues 24–153, 190–239, 276–318 and 361–394; the GCGR input contained residues 26–154, 183–240, 273–316 and 359–394; and the GIPR input contained residues 31–148, 179–232, 264–308 and 351–383. Pairwise joint-target campaigns were performed for GLP-1R–GCGR, GLP-1R–GIPR and GCGR–GIPR and compared with corresponding campaigns in which peptides were optimised against each receptor individually. The same cropped target structures, binding-site definitions and remaining design settings were used for the joint- and single-target comparisons. The campaign-specific optimisation schedule is summarised in Table 1. The semigreedy stage evaluated discrete mutations without gradient-based sequence updates. Three single-target campaigns were performed, one for each receptor. Each single-target campaign was reused as the control in the two pairwise comparisons containing its optimised receptor. Primary-analysis cohort sizes are reported in Table 1. Independent GPCR re-evaluation was performed using AF2-Multimer rather than AF3 because AF3 was not available for this benchmark under the access and licensing terms applicable when the analysis was conducted. To provide an independent prediction setup, AF2-Monomer-predicted receptor structures were supplied to AF2-Multimer as initial-guess coordinates during re-prediction of the peptide–receptor complexes. This target-preparation strategy was selected because related AF2-guided peptide-binder design campaigns have produced experimentally validated binders using AF2-predicted target structures [15]. The re-evaluation was intended to test whether the joint-optimisation signal persisted under an independent AF2 prediction setup, rather than to provide an AF3 comparison.

#### 4.2.3 Three-finger toxin cross-reactivity benchmark

The 3FTx benchmark compared minibinders optimised against Erabutoxin A alone with minibinders jointly optimised against Erabutoxin A and NK-shNTx. Both conditions used the same primary-target structure and remaining optimisation settings. Cohort sizes are reported in Table 1.

#### 4.2.4 pMHC M4A specificity benchmark

The pMHC benchmark used SLLMWITQC/HLA-A*02:01 as the on-target and the single-residue M4A variant, SLLAWITQC/HLA-A*02:01, as the off-target. Designs generated with an M4A off-target objective were compared with designs generated without off-target counter-selection, while the remaining optimisation settings were held fixed. The off-target context weight is reported in Table 1. For the off-target, clipped normalised iPAE and iPTM loss terms were applied. Normalised iPAE was defined as the raw iPAE divided by 31 Å, yielding a dimensionless value on an approximately 0–1 scale. The normalised-iPAE loss was active only when the value was below 0.35, whereas the iPTM loss was active only when iPTM exceeded 0.45; otherwise, the corresponding loss term was set to zero. The off-target hotspot-contact loss penalised, for each binder residue, predicted contact probabilities with the specified off-target hotspot residues above 0.30, using a 20 Å distance cutoff and a weight of 0.05. The off-target objective was combined with the on-target objective using the RMS-normalised PCGrad procedure defined under *Gradient normalisation and combination*.

The campaign-specific optimisation schedule, including the modified final straight-through one-hot stage, is summarised in Table 1.

After retaining designs with complete trajectory records, cohorts were capped chronologically by completion time, and the same design sets were used for the AF2 and AF3 analyses. Cohort sizes are reported in Table 1.

### 4.3 Three-finger toxin minibinder design and experimental validation

#### 4.3.1 Computational design and selection

Cross-reactive 80-residue minibinders were designed against Erabutoxin A from *Laticauda semifasciata* and NK-shNTx from *Naja kaouthia* (UniProt ID P59276) using Odin-Multi. Each toxin was evaluated in a separate AF2-Multimer minibinder-design task while sharing the same minibinder sequence representation. Both toxins were assigned target roles and jointly contributed to optimisation of the shared minibinder sequence. Campaign-specific context weights, ProteinMPNN regularisation and optimisation settings are summarised in Table 1. The per-target structural objective followed the on-target loss function defined under *On-target and off-target loss functions*, and cross-reactivity candidates were ranked using *S*_cross_ as defined under *Candidate ranking*.

Two design batches were performed. The first batch used no hotspot constraints, whereas the second specified residues 51–55 of Erabutoxin A and the alignment-equivalent residues 50–54 of NK-shNTx as binding-site hotspots. Trajectory counts are reported in Table 1. Each two-target trajectory took a median of 23.3 min on a T4. Poly 5 was generated in the second, hotspot-constrained batch. For each trajectory, the AF2 candidates selected according to the procedure summarised in Table 1 were modelled as separate binary complexes with each toxin using AF3. Predictions were evaluated using iPAE, ipSAE, minibinder pLDDT, and Rosetta interface shape complementarity. The final campaign-specific filtering thresholds are reported in Table 1. A final panel of 30 designs satisfying these joint-target criteria was selected for experimental testing.

#### 4.3.2 Cloning and protein production

Minibinder sequences were codon optimised for *E. coli* expression using GenScript’s online tool. BsaI cut sites and plasmid-compatible overhangs were added for Golden Gate cloning [42] and the constructs were ordered from GenScript. Cloning, expression, and purification were performed in 96-well formats using a workflow adapted from Qian *et al.* [43]. The constructs were cloned into a custom pETmod24a+-based vector (GenScript) containing a start-codon, a ramp sequence at codon 3-4 (AACATT) [44], and a C-terminal 3xFLAG-tag and C-tag separated by GS-based linkers for detection and purification purposes. The plasmids were transformed into CaCl_2_-based chemically competent BL21(DE3) cells, plated out, and single colonies chosen for inoculation of 4 mL of autoinduction media per construct [45]. Cells were grown overnight, pelleted using centrifugation, and lysed in 50 mM TRIS (pH 7.4), 0.1% Triton X-100, 1 mM MgCl_2_, 0.1 mg/mL lysozyme (Sigma Aldrich) and 0.05 *µ*L/mL benzonase (EMD Millipore Corp.). Lysate was pelleted and the supernatant transferred to 50 *µ*L CaptureSelect^TM^ C-tagXL resin (ThermoFisher Scientific) in a fritted plate placed on a NucleoVac 96 Vacuum Manifold (Takara). The resin was washed with TRIS buffer saline (TBS pH 7.4, 50 mM TRIS, 150 mM NaCl) and protein eluted using 180 *µ*L 0.1 M glycine (pH 2.7) into 20 *µ*L neutralisation buffer (1 M TRIS pH 8, 1.5 M NaCl) for a final pH of 7.4, thereby omitting the need of buffer exchange for our subsequent applications. The protein was quantified in a 384-well plate format using a Qubit protein quantification kit (ThermoFisher Scientific) and protein concentrations normalised to 3 *µ*M. Constructs that yielded less than 3 *µ*M were not diluted further (ranging 1.7 to 2.5 *µ*M).

#### 4.3.3 Minibinder screening using DELFIA

Minibinders were screened using dissociation-enhanced lanthanide fluorescence immunoassay (DELFIA). MaxiSorp plates (ThermoFisher Scientific) were coated overnight in PBS with 2 *µ*g/mL purified Erabutoxin A from *L. semifasciata*, purified *N. pallida* short neurotoxin 1 (NP-shNTx), or *N. kaouthia* whole venom and were subsequently blocked with PBS containing 0.1% Tween-20 and 1% skimmed milk. NP-shNTx was included as the closest available purified homologue of NK-shNTx because purified NK-shNTx was unavailable. A block-only control was included to assess non-specific binding. Minibinders were incubated at 200 nM in blocking buffer, followed by Europium-labelled anti-FLAG M2 antibody (F3165, Sigma-Aldrich) in DELFIA assay buffer (Perkin Elmer). The plate was developed using DELFIA enhancement solution (PerkinElmer), and fluorescence was measured at 337 nm excitation and 615 nm emission using a VICTOR Nivo plate reader (PerkinElmer). Target-associated binding was evaluated by comparing the signal from each toxin-containing coating with the block-only control.

#### 4.3.4 Large-scale production of hit minibinder

The plasmid encoding the hit minibinder Poly 5 was confirmed by sequencing (Eurofins Genomics). A 2xYT culture was inoculated with the clone, induced with IPTG, and incubated overnight at 20 *^◦^*C. The culture was spun down, the pellet resuspended in TBS (pH 7.4), 0.05 *µ*L/mL benzonase, 1 mM MgCl_2_ and cOmplete EDTA-free protease inhibitors (Roche), and lysed using sonication. The lysate was spun down, the supernatant filtered on a 0.45 *µ*m syringe filter, and purified on an NGC BioRad system using a CaptureSelect^TM^ C-tagXL prepacked 5 mL column (ThermoFisher Scientific). TBS served as wash buffer and 2 M MgCl_2_ as elution buffer. The column was stripped with 0.1 M glycine (pH 2). The protein was concentrated on a 10 kDa MWCO Amicon centrifugal filter (Sigma Aldrich) and applied for size exclusion chromatography (SEC) in TBS using a Superdex 75 10/300GL column (Cytiva) on an NGC BioRad system. The size of the minibinder was confirmed using SDS-PAGE.

#### 4.3.5 Determining binding kinetics using bio-layer interferometry

Binding kinetics were determined using an Octet RED 96 (Fortebio) using Octet Discovery (12.2.2.20) and Octet Analysis (12.2.2.4). All assays were performed in 10 mM HEPES, 150 mM NaCl, 3 mM EDTA, 50 mM MES hydrate, and 0.05% P_20_. Erabutoxin A and venom-derived fraction 3 were biotinylated using ChromaLINK (Vector Laboratories) and loaded onto streptavidin biosensors (Sartorius) to a loading response of approximately 1 nm. Poly 5 was tested at six twofold serial dilutions from 400 to 12.5 nM. Target against buffer only was subtracted from the signals. The curves were fitted using a 2:1 heterogeneous-ligand model.

#### 4.3.6 RP-HPLC fractionation and MALDI-TOF analysis

Five milligrams of *Naja kaouthia* whole venom (Latoxan, Cat.# L1323) was reconstituted in water containing 0.1% trifluoroacetic acid (TFA). The venom was fractionated by reversed-phase high-performance liquid chromatography (RP-HPLC) on an Agilent 1200 system equipped with a C18 column (250 *×* 4.6 mm, 5 *µ*m particle size; Teknokroma). Elution was performed at a flow rate of 3 mL/min using buffer A (0.1% TFA in water) and buffer B (0.1% TFA in acetonitrile). Buffer B was increased from 0 to 5% over 10 min, from 5 to 15% over the following 20 min, and from 15 to 30% over the subsequent 120 min. Four fractions were collected, and the solvent was removed using a vacuum centrifuge. The dried fractions were reconstituted in water and analysed by matrix-assisted laser desorption/ionisation time-of-flight (MALDI-TOF) mass spectrometry using a ProteoMass Protein MALDI-MS Calibration Kit (Sigma-Aldrich, Cat.# MSCAL3) and an Autoflex Speed MALDI/TOF mass spectrometer (Bruker Daltonics).

Candidate identities were informed by a previously published *N. kaouthia* venomics analysis, which identified the closely related short-chain neurotoxins P59275 (6944 Da) and P59276 (6859 Da), together with the more distantly related 3FTx homologue E2IU03, within a mixed RP-HPLC peak [32]. Fraction 3 showed a dominant mass of 6894.743 Da, falling between the theoretical masses of P59276 and P59275, and was therefore selected as the candidate NK-shNTx-containing fraction for subsequent binding analysis. The imperfect mass match could reflect sequence variation, including an amino-acid substitution, although intact-mass MALDI-TOF alone cannot establish this.

### 4.4 pMHC minibinder design and experimental validation

#### 4.4.1 Computational design and selection

Eighty-residue minibinders were designed against SLLMWITQC/HLA-A*02:01 under three conditions. The target-only condition included only the on-target and did not include an explicit off-target objective. The CMV counter-selection condition included NLVPMVATV/HLA-A*02:01 and empty HLA-A*02:01 as off-targets. The M4A counter-selection condition included SLLAWITQC/HLA-A*02:01 and empty HLA-A*02:01 as off-targets. Campaign-specific context weights are reported in Table 1.

Per-trajectory runtime was determined by the number of prediction contexts and by GPU type. The single-context target-only condition took a median of 20.2 min per trajectory on a V100. The three-context counter-selected conditions took 61.5 min (CMV) and 59.3 min (M4A) per trajectory on a V100, and 158.7 min and 158.1 min, respectively, on a T4. Adding the two off-target contexts therefore increased per-trajectory cost approximately threefold at matched GPU type, and the T4 was approximately 2.6-fold slower than the V100 for the same objective.

Candidates were re-evaluated and filtered using the AF3 procedure defined under *Filtering*, with the final campaign-specific criteria reported in Table 1. The final library contained 166 designs from the target-only condition and 167 designs from each of the CMV- and M4A-counter-selected conditions, yielding a pooled library of 500 designs.

#### 4.4.2 Recombinant expression of MHC molecules and generation of peptide MHC complexes

The MHC molecules were expressed as previously described [46]. Briefly, recombinant HLA-A*02:01 heavy chain, with a Y84C mutation, and a light chain of human *β*_2_ microglobulin (*β*_2_*M*) were produced in chemically competent *E. coli* from One Shot^TM^ BL21(DE3) pLysS (Invitrogen, Cat.# C606003). Heavy and light chains were refolded with 10 *µ*M dipeptide GM. Resulting monomers were biotinylated and purified by size exclusion chromatography using high-performance liquid chromatography. For generating peptide-HLA-A*02:01-Y84C complexes, purified HLA-A*02:01-Y84C monomers (200 *µ*g/mL) were incubated with 400 *µ*M peptide (SB-PEPTIDE, France) in phosphate-buffered saline (PBS) for 30 min at RT. The resulting peptide MHC (pMHC) complexes were centrifuged for 5 min at 10,000 *×g*, and supernatants were used to generate pMHC tetramers.

#### 4.4.3 Structural analysis of the S2 complex

The AF3-predicted S2 complex was analysed to examine the environment of peptide Met4 (Fig. **4**f). Coordinates were used without further relaxation or energy minimisation. Contacts were defined by the minimum Euclidean distance between any Met4 side-chain heavy atom (C*β*, C*γ*, S*δ*, or C*ε*) and any heavy atom of each minibinder residue, including backbone atoms. Residues with a minimum distance *≤* 4.2 Å were included, and the closest atom pair per residue was displayed with distances rounded to 0.1 Å. This identified Met20, Val23, Met54, Phe57, Glu58, and Trp61; the Met54 contact involved its backbone carbonyl oxygen. These contacts describe geometric proximity without assigning hydrogen bonds or interaction energies. The complex overview was rendered using PyMOL version 3.0.0, and pocket and contact close-ups using Mol* version 4.9.0. Distance calculations, annotations, and figure assembly used custom Python scripts.

#### 4.4.4 Preparation of pMHC tetramers

For each 100 *µ*L of pMHC complexes, 9 *µ*L of streptavidin conjugates [0.2 mg/mL stock SAPE (Biolegend, Cat.#405204) or SA-APC (Biolegend, Cat.#405243)] was added and incubated for 30 min at 4 *^◦^*C, followed by the addition of D-biotin (Sigma-Aldrich, Cat.#2031) at a final concentration of 25 *µ*M to block any free binding sites. Assembled pMHC tetramers were stored at −20 *^◦^*C with 5% glycerol and 0.5% BSA.

#### 4.4.5 Maintenance of cell lines

HEK293T cells (ATCC, Cat.#CRL-3216, RRID: CVCL 0063) and Jurkat cell lines (Signosis, Cat.#SL-0032) were cultured in complete R10 (RPMI 1640 + GlutaMAX^TM^-I (Gibco, Cat.#61870-010) with 10 % FBS (Biowest, Cat.#S181H), 1% Pen-Strep (Merck Life Science A/S, Cat.#P0781)) at 37 *^◦^*C, 5% CO_2_, and split every 2–4 days before confluence. Jurkat T CD3 KO cells were produced as described in [18].

#### 4.4.6 Gene design and cloning

A library of 500 miBd designs against SLLMWITQC/HLA-A*02:01 was synthesised as an oligo pool (Twist Bioscience). The library was cloned into the CAR transfer vector (pMB-03), a modified version of pLenti-puro (Addgene, Cat.#39481, RRID: Addgene 39481), optimised to include a cPPT-CTS sequence, the EF-1*α* promoter, a Woodchuck Hepatitis Virus (WHV) Post-transcriptional Regulatory Element (WPRE), a CD8 leader sequence, BsmBI cloning sites, the CAR domains CD8*α* hinge-41BB-CD3*ζ*, and truncated nerve growth factor receptor (tNGFR). Cloning of miBds into pMB-03 was performed using NEBridge^®^ Golden Gate Assembly Kit (BsmBI v2) (NEB, Cat.#E1602) according to manufacturer’s protocol. Golden Gate products were electroporated into ElectroMAX STBL4 competent bacteria (Invitrogen, Cat.# 11635018) with the program ‘*E. coli* – 1mm, 1.8 kV’ on Gene Pulser Xcell Electroporation System (Bio-Rad, Cat.#1652660). Library coverage was assessed by PCR using NEBNext^®^ Ultra^TM^ II Q5^®^ Master Mix (NEB, Cat.#M0544) and primers MB GGinterm fwd 2 (TTCATTCTCAAGCCTCA-GACAGT) and MB GGinterm rev 2 (GATATCGCAGGCGAAGTCAAGA). PCR amplified miBd sequences were labelled with Native Barcoding Kit 96 V14 (Nanopore, Cat.#SQK-NBD114.96) and sequenced by Nanopore PromethION. The resulting FASTQ reads were analysed using a custom Python script that matched the sequences against the minibinder sequence library.

#### 4.4.7 Mammalian surface display of miBds

Library screening was performed as previously described in [18]. Briefly, lentiviral particles containing the minibinder library were produced by lipofectamine-based co-transfection of HEK293T cells with 3rd generation packaging plasmids pMD2.G (Addgene, Cat.#12259), pMDLg/pRRE (Addgene, Cat.#12251), pRSV-Rev (Addgene, Cat.#12253), and the pMB-03 plasmid with the library of minibinder sequences. Lentiviral supernatant was collected 24 and 48 hours after transfection, pooled, concentrated (Lenti-X^TM^ Concentrator, Takara Bio, Cat.#631232), and stored at −80 *^◦^*C. CD3 KO Jurkat T cells were transduced with concentrated lentivirus across a series of viral dilutions. Transduction efficiency was assessed by anti-tNGFR staining (clone C40–1457, BD, Cat.# 743362, RRID: AB 2741453). A dilution resulting in approximately 40% tNGFR-positive cells was selected to favour single viral integration events per cell. For the library transduction, 2 *×* 10^6^ CD3 KO Jurkat cells were used. Tetramers (SLLMWITQC/HLA-A*02:01) were separately labelled with PE and APC fluorochromes and pooled before cell staining. The CD3 KO Jurkat cells containing the library of miBds were stained with LIVE/DEAD^TM^ Fixable Near-IR Dead Cell Stain Kit (ThermoFisher, Cat.#L10119) and anti-tNGFR to evaluate minibinder expression. Binding to the pMHC target was evaluated by double PE- and APC-tetramer staining. Cells were sorted into Live, tNGFR^+^, PE/APC double-positive and PE/APC double-negative populations on a BD FACSDiscover^TM^ S8 Cell Sorter (BD, RRID: SCR 026674). Genomic DNA was extracted using QIAamp DNA Micro Kit (Qiagen, Cat.#56304) following the manufacturer’s instructions. To amplify the integrated miBd DNA from the integrated CAR construct, PCR amplification and Nanopore sequencing were performed using the workflow described under *Gene design and cloning*. FASTQ read files were analysed using the custom sequence-matching Python workflow described under *Gene design and cloning*. For each sample, miBd read counts were converted to relative abundances by dividing each count by the total number of reads in that sample. Log_2_ fold changes (log_2_FC) were calculated between sorted populations using the normalised read fractions. The 20 designs with the highest log_2_ FC values across the pooled library were selected for individual clone characterisation; the lowest-ranked selected design had a log_2_ FC of approximately 1.2.

#### 4.4.8 Cross-reactivity assessment using pMHC tetramers

The identified miBd sequences able to bind SLLMWITQC/HLA-A*02:01 were cloned as single constructs and transduced into CD3 KO Jurkat cells using the lentiviral transduction procedure described under *Mammalian surface display of miBds*. Target binding was confirmed using PE-labelled SLLMWITQC/HLA-A*02:01 tetramers. Cross-reactivity was evaluated using PE-labelled tetramers presenting the single-residue M4A variant SLLAWITQC/HLA-A*02:01 and the sequence-divergent CMV peptide NLVPMVATV/HLA-A*02:01. Following tetramer staining, cells were stained with anti-tNGFR and LIVE/DEAD^TM^ Fixable Near-IR using the reagents and staining procedure described under *Mammalian surface display of miBds*. Data were acquired using an LSRFortessa^TM^ flow cytometer and analysed using FlowJo. Cells were gated on live singlets followed by selection of tNGFR^+^ cells to confirm miBd expression.

### 4.5 Statistical analysis

Fixed-threshold pass-rate comparisons between optimisation conditions in the GPCR and 3FTx benchmarks were evaluated using two-sided Fisher’s exact tests on the corresponding 2 *×* 2 count tables.

Because no single target-to-off-target iPTM-ratio threshold was pre-specified for the pMHC benchmark, the primary analysis compared the complete target-qualified specificity-yield curves. For each of 160 evenly spaced ratio thresholds from 1 to 5, yield was calculated as the fraction of all designs with target iPTM *>* 0.5 and a target-to-off-target iPTM ratio exceeding the evaluated threshold. The global test statistic was the maximum absolute difference in yield between the two conditions across this threshold range. Its significance was assessed by permuting the design-condition labels 50,000 times using seed 20260804, while retaining each design’s paired target and off-target scores and the original condition sizes. The same curve-comparison procedure was applied to the corresponding AF3 re-evaluations. Results at ratio thresholds of 1.5, 2.0, 2.5 and 3.0 were reported descriptively without separate hypothesis tests; the threshold of 2.5 was selected post hoc and was not used as the primary statistical comparison.

Holm adjustment was applied separately to three families of comparisons: the eight primary AF2 comparisons, including the pMHC permutation *P* value; the six GPCR AF2 re-evaluation comparisons; and the two AF3 re-evaluation comparisons. All adjusted values are reported in Supplementary Table 1.

Pointwise uncertainty for the threshold-yield curves shown in Fig. **2**b–d was estimated independently for each condition using 2,000 bootstrap resamples. Each bootstrap replicate retained the original condition-specific sample size, and yield was recalculated at every evaluated threshold. The resulting empirical distributions were used to calculate pointwise 95% bootstrap percentile intervals. These intervals describe uncertainty at individual thresholds and were not interpreted as separate hypothesis tests. Designs were not pooled between campaigns or optimisation conditions.

## Data Availability

The data supporting the findings of this study are provided within the manuscript and its Supplementary Information. Additional source data are available on Zenodo at https://doi.org/10.5281/zenodo.22292558.

## Code Availability

The source code for Odin-Multi is available at https://github.com/DigBioLab/odin_multi.

## Author Contributions

V.B. and T.P.J. conceived the study. V.B. developed Odin-Multi and performed the computational design campaigns and analyses. C.R.C. and B.S. performed the pMHC experiments; K.H.B. and M.B.-V. performed the snake-venom experiments; V.E.M. performed the GPCR benchmarks. M.M.N., J.C.N., S.R.H., K.H.J. and T.P.J. supervised the work. V.B. and T.P.J. wrote the manuscript, with experimental sections drafted by C.R.C. and K.H.B. All authors reviewed and approved the final manuscript.

## 5 Supplementary materials

**Supplementary Figure 1:**
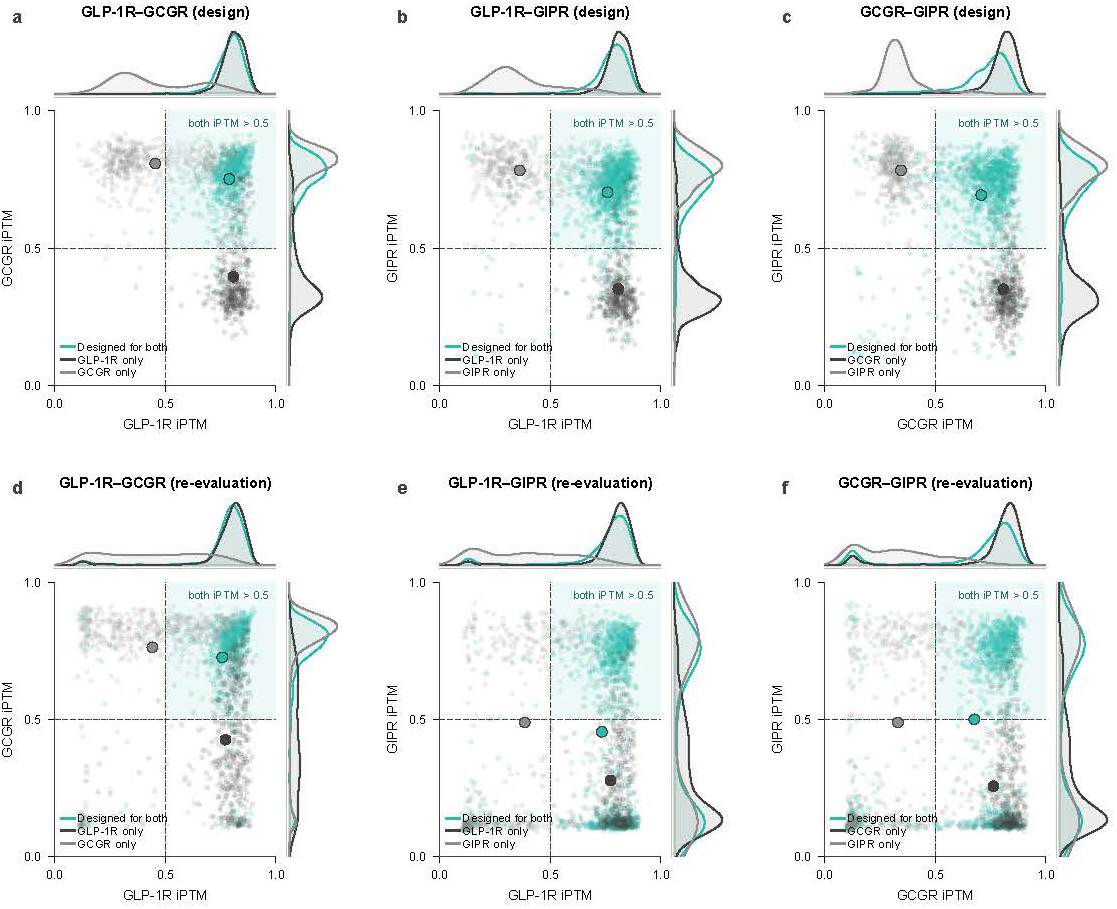
Pairwise GPCR interaction-confidence distributions before and after independent AF2 re-evaluation. **a–c**, Original pairwise iPTM distributions for designs targeting GLP-1R and GCGR (**a**), GLP-1R and GIPR (**b**), or GCGR and GIPR (**c**). **d–f,** Corresponding iPTM distributions after each design was independently re-evaluated against GLP-1R, GCGR, and GIPR (**d–f**). Teal denotes designs jointly optimised against both receptors, whereas black and grey denote the corresponding single-receptor campaigns, as identified in each panel. Individual points represent designs, enlarged circles indicate group means, and the top and right marginal plots show kernel-density estimates for the respective receptor scores. Dashed lines indicate an iPTM threshold of 0.5, and the shaded upper-right quadrant denotes designs exceeding this threshold for both receptors. The primary design-time distributions use the final optimisation iteration, whereas the independent re-evaluation uses the sequence retained by the original minimum-loss rule for each GPCR trajectory. Cohort sizes differ slightly between design and re-evaluation because each panel uses every design for which that measurement exists: eight GLP-1R–GIPR joint designs have no re-evaluation record, and 35 GLP-1R-only designs have no complete trajectory record.

**Supplementary Figure 2:**
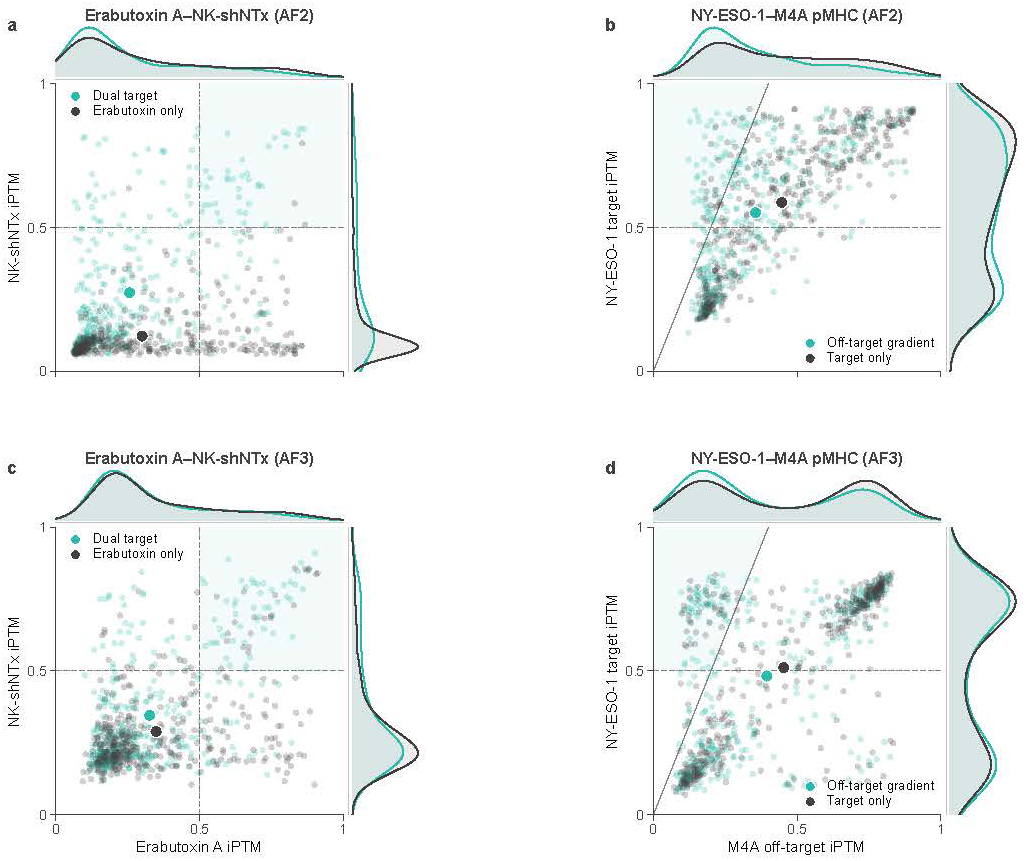
Joint distributions of predicted interaction confidence for cross-reactive and specificity-optimised Odin-Multi designs. **a**, AlphaFold2-Multimer iPTM scores for Erabutoxin A and NK-shNTx. Designs jointly optimised against both toxins are compared with designs optimised against Erabutoxin A alone. Dashed lines indicate an iPTM threshold of 0.5 for each target, and the shaded upper-right quadrant denotes designs exceeding this threshold for both toxins. **b,** AlphaFold2-Multimer iPTM scores for the NY-ESO-1 target pMHC, SLLMWITQC/HLA-A*02:01, and the M4A off-target pMHC, SLLAWITQC/HLA-A*02:01. Designs generated with an explicit off-target gradient are compared with designs generated without off-target counter-selection. The horizontal dashed line indicates the target iPTM threshold of 0.5 used to define target-qualified yield, whereas the diagonal line indicates the post hoc illustrative target-to-off-target iPTM ratio threshold of 2.5. The primary pMHC analysis compared the complete specificity-yield curves over ratio thresholds from 1 to 5 rather than testing this single ratio cutoff. **c,** Corresponding AlphaFold3 predictions for the Erabutoxin A and NK-shNTx design sets. **d,** Corresponding AlphaFold3 predictions for the NY-ESO-1 target and M4A off-target design sets, with the same dashed thresholds shown for descriptive reference. In all panels, individual points represent designs, enlarged circles indicate group means, and marginal density plots show the corresponding iPTM distributions. Teal denotes joint-target optimisation or optimisation with an off-target gradient, whereas grey denotes the corresponding single-target or no-off-target-gradient control.

**Supplementary Figure 3:** Expression and biochemical characterisation of Odin-Multi-designed cross-reactive toxin minibinders. **a**, Protein yields following small-scale expression and purification of 30 Odin-Multi-designed minibinders. An *α*-cobratoxin-targeting miBd (acbtx miBd) and an empty-plasmid negative control (Neg. con.) were included as controls. The dashed line indicates the yield of the negative control (6.9 *µ*g); 24 of the 30 designs produced yields above this value. **b,** RP-HPLC separation of *Naja kaouthia* whole venom monitored by absorbance at 215 nm. The solid line shows the chromatographic absorbance profile, the dashed line shows the acetonitrile gradient, and the shaded regions indicate the collection windows for fractions F1–F4. **c,** SDS-PAGE analysis of the four collected RP-HPLC fractions. M denotes the molecular-weight marker, F1–F4 denote the collected fractions, and Era denotes purified Erabutoxin A included as a reference. **d,** SDS-PAGE analysis of purified Poly 5 following large-scale expression and purification, showing a predominant band at the expected molecular mass of 14.7 kDa.

**Supplementary Figure 4:**
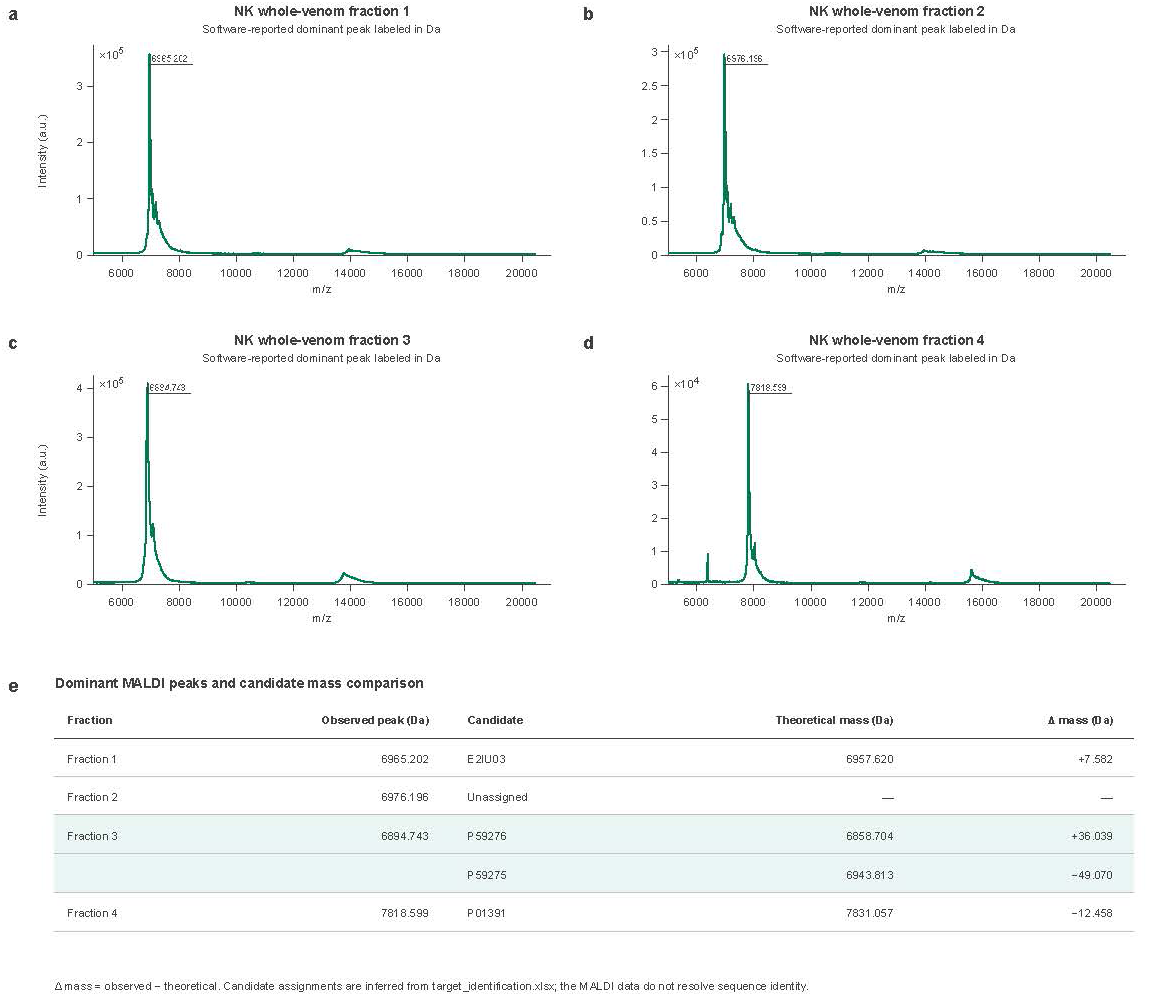
MALDI-TOF mass analysis of RP-HPLC fractions isolated from *Naja kaouthia* whole venom. **a**, MALDI-TOF spectrum of fraction 1, with a software-reported dominant peak at 6965.202 Da. **b,** MALDI-TOF spectrum of fraction 2, with a software-reported dominant peak at 6976.196 Da. **c,** MALDI-TOF spectrum of fraction 3, with a software-reported dominant peak at 6894.743 Da. **d,** MALDI-TOF spectrum of fraction 4, with a software-reported dominant peak at 7818.599 Da. **e,** Comparison of the observed dominant masses with the theoretical masses of candidate venom proteins. Fraction 1 was compared with E2IU03, fraction 3 with the short-chain neurotoxin candidates P59276 and P59275, and fraction 4 with P01391; fraction 2 remained unassigned. Δ mass was calculated as the observed mass minus the theoretical mass. Candidate assignments were inferred from mass proximity and should therefore be regarded as tentative, because MALDI-TOF mass measurements alone do not establish sequence identity.

**Supplementary Figure 5:**
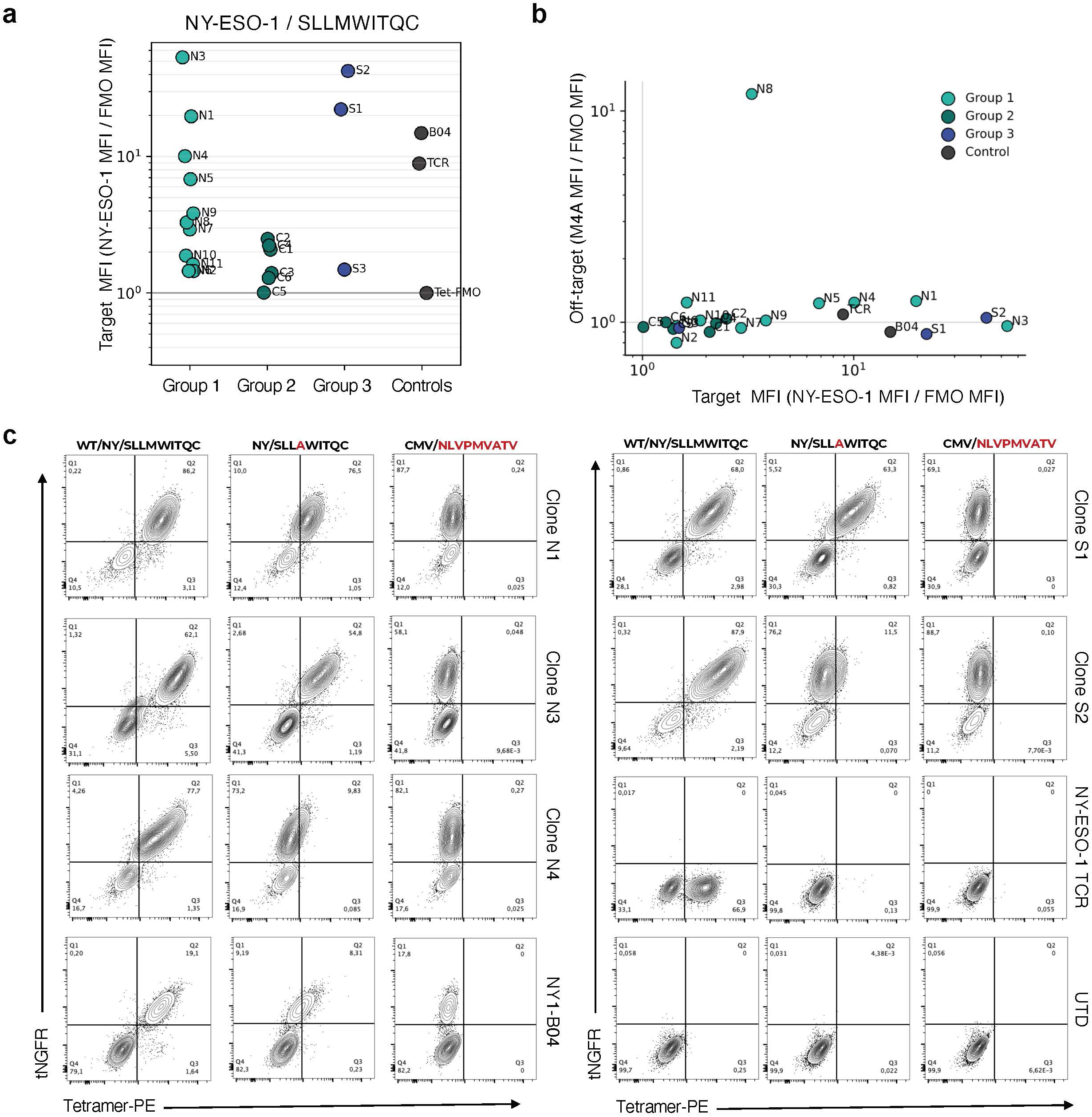
Experimental validation of pMHC-targeting miBds designed using Odin-Multi. **a**, Dot plot showing PE-labelled NY-ESO-1 target tetramer staining of the 20 selected individual clones. Designs are grouped by design condition: Target-only (Group 1), CMV-counter-selected (Group 2), and SLLAWITQC (M4A)-counter-selected (Group 3), in which binding to the single-residue SLLAWITQC variant was penalised during design. MFI values were normalised to the tetramer FMO control, with values above 1 indicating enrichment over the background. Cells were gated on singlet/live/tNGFR^+^ cells. **b,** Dot plots showing individual tetramer staining of the 20 selected clones with the NY-ESO-1 target tetramer and the CMV off-target tetramer (NLVPMVATV/HLA-A*02:01). MFI values were similarly normalised to the corresponding tetramer FMO control. **c,** Flow plots of selected clones stained with PE-labelled tetramers presenting the NY-ESO-1 target peptide, the M4A off-target peptide, or the CMV off-target peptide. Shown are three clones from the target-only design condition (N1, N3, N4), two clones from the M4A-counter-selected group (S1, S2), the NY1-B04, the NY-ESO-1-specific TCR control (1G4), and untransduced CD3 KO Jurkat cells. Cells were gated on singlet/live cells. All miBd clones were expressed as 41BB–CD3*ζ* CARs in CD3 KO Jurkat cells. The NY-ESO-1-specific TCR control (1G4) was expressed as a TCR and did not contain the tNGFR transduction marker. Tet-FMO denotes the tetramer fluorescence-minus-one control, and UTD denotes the no-binder control comprising untransduced CD3 KO Jurkat cells.

---

**Supplementary Table 1.** Statistical comparisons of computational benchmark performance. Fixed-threshold binary comparisons were evaluated using two-sided Fisher’s exact tests. The pMHC comparisons used global permutation tests of the complete target-qualified specificity-yield curves. *P* values were Holm-adjusted separately across the eight primary AF2 comparisons, the six GPCR AF2 re-evaluation comparisons, and the two AF3 re-evaluation comparisons.

| System | Evaluation | Odin-Multi condition | Control condition | Holm-adjusted $P$ |
| --- | --- | --- | --- | --- |
| GLP-1R–GCGR | AF2 design | Dual: 596/616 (96.8%) | GLP-1R: 74/393 (18.8%) | $1.27 \times 10^{-159}$ |
| GLP-1R–GCGR | AF2 design | Dual: 596/616 (96.8%) | GCGR: 168/463 (36.3%) | $4.47 \times 10^{-114}$ |
| GLP-1R–GIPR | AF2 design | Dual: 999/1,102 (90.7%) | GLP-1R: 42/393 (10.7%) | $9.55 \times 10^{-193}$ |
| GLP-1R–GIPR | AF2 design | Dual: 999/1,102 (90.7%) | GIPR: 71/381 (18.6%) | $5.97 \times 10^{-154}$ |
| GCGR–GIPR | AF2 design | Dual: 921/1,103 (83.5%) | GCGR: 49/463 (10.6%) | $2.82 \times 10^{-170}$ |
| GCGR–GIPR | AF2 design | Dual: 921/1,103 (83.5%) | GIPR: 26/381 (6.8%) | $6.56 \times 10^{-167}$ |
| 3FTx | AF2 design | Dual: 37/400 (9.2%) | Erabutoxin-only: 3/400 (0.8%) | $1.86 \times 10^{-8}$ |
| pMHC | AF2 design | Off-target gradient: 57/400 (14.2%) | No off-target gradient: 24/400 (6.0%) | $7.00 \times 10^{-4}$ |
| GLP-1R–GCGR | AF2 re-eval. | Dual: 539/616 (87.5%) | GLP-1R: 161/428 (37.6%) | $1.70 \times 10^{-64}$ |
| GLP-1R–GCGR | AF2 re-eval. | Dual: 539/616 (87.5%) | GCGR: 190/463 (41.0%) | $1.52 \times 10^{-59}$ |
| GLP-1R–GIPR | AF2 re-eval. | Dual: 548/1,094 (50.1%) | GLP-1R: 59/428 (13.8%) | $4.65 \times 10^{-42}$ |
| GLP-1R–GIPR | AF2 re-eval. | Dual: 548/1,094 (50.1%) | GIPR: 87/381 (22.8%) | $2.79 \times 10^{-21}$ |
| GCGR–GIPR | AF2 re-eval. | Dual: 599/1,103 (54.3%) | GCGR: 52/463 (11.2%) | $7.97 \times 10^{-62}$ |
| GCGR–GIPR | AF2 re-eval. | Dual: 599/1,103 (54.3%) | GIPR: 52/381 (13.6%) | $9.92 \times 10^{-47}$ |
| 3FTx | AF3 re-eval. | Dual: 50/400 (12.5%) | Erabutoxin-only: 22/400 (5.5%) | 0.0015 |
| pMHC | AF3 re-eval. | Off-target gradient: 59/400 (14.8%) | No off-target gradient: 38/400 (9.5%) | 0.0612 |

*Note:* GPCR AF2 design rows use the final optimisation iteration, applied identically to dual and single-target campaigns, so no pair-aware post-hoc selection is applied to the controls. The independent GPCR AF2 re-evaluation used the exact design-ID cohorts available in the re-evaluation dataset. Consequently, its cohort sizes differ from those of the primary fixed-checkpoint analysis: the GLP-1R-only cohort contains 428 rather than 393 designs, and eight GLP-1R–GIPR dual designs lacked re-evaluation results. The AF3 rows use the native-target-MSA re-evaluation of the exact final stored trajectory sequence (seed 1, mean over five samples), with no post-hoc iteration selection.

**Supplementary Table 2.** Global kinetic parameters from Octet 2:1 heterogeneous-ligand fits with independent Rmax for Poly 5 binding to Erabutoxin A and candidate NK-shNTx-containing fraction 3. Values are best-fit estimates *±* Octet-reported standard errors.

| Target | Fit component | $K_D$ (nM) | $k_{on}$ ($M^{-1} s^{-1}$) | $k_{off}$ ($s^{-1}$) | $R^2$ |
| --- | --- | --- | --- | --- | --- |
| Erabutoxin A | 1 (tight) | $11.95 \pm 0.22$ | $(3.23 \pm 0.02) \times 10^4$ | $(3.86 \pm 0.07) \times 10^{-4}$ | 0.9959 |
| Erabutoxin A | 2 (weak) | $48.41 \pm 0.92$ | $(2.16 \pm 0.03) \times 10^5$ | $(1.05 \pm 0.01) \times 10^{-2}$ | 0.9959 |
| NK whole-venom fraction 3 | 1 (tight) | $34.43 \pm 0.26$ | $(4.35 \pm 0.03) \times 10^4$ | $(1.50 \pm 0.01) \times 10^{-3}$ | 0.9984 |
| NK whole-venom fraction 3 | 2 (weak) | $77.49 \pm 1.07$ | $(2.27 \pm 0.03) \times 10^5$ | $(1.76 \pm 0.01) \times 10^{-2}$ | 0.9984 |

*Note:* “Tight” and “weak” denote the two fitted kinetic components of the heterogeneous-ligand model. Fraction 3 is a venom-derived fraction whose short-chain three-finger toxin identity was inferred from chromatographic behaviour and intact-mass comparison. The fitted components are not assigned individually to the proteins present in the fraction. The reported $R^2$ applies to the corresponding global fit and is therefore repeated for both components.

